# Dopamine signaling tracks naturalistic short-term fluctuations in hunger-satiety

**DOI:** 10.1101/2025.04.29.651295

**Authors:** Devry Mourra, Cayla E Murphy, Elias Hourany, Kelly Centeno, Federico Gnazzo, Jeff A Beeler

## Abstract

The neuromodulator dopamine is integral to feeding behavior, believed to modulate food pursuit and satiety. Here, we examine how dopamine signaling in the nucleus accumbens changes during consumption as animals transition from hunger to satiety in naturalistic feeding. Dopamine transiently increases during food approach; however, the magnitude of this approach-related increase diminishes across progressive pellet ingestion, reflecting short-term satiation. These approach-related dopamine transients recover during intermeal intervals. Fasting dissociates the regulation of meal size and frequency, reflecting termination and initiation, respectively, with observable differences in dopamine corresponding to changes in meal size but not frequency. Despite substantially decreasing feeding, pharmacological satiation via a glucagon-like peptide-1 (GLP-1) agonist had no impact on average dopamine transients on food approach but abolished the short-term fluctuations related to on-going eating, suggesting that the GLP-1 agonist disengaged or decoupled dopamine from its modulatory role in meal patterning.

## Introduction

Critical to energy management, appetite and satiation are highly regulated, complex and contain many redundant systems (1,2). A negative energy balance can lead to malnutrition and death, while a surplus increases the risk of obesity (3). While motivation for food pursuit is critical, limiting consumption is equally necessary.

Dopamine (DA) is widely believed to play a central role in driving appetitive pursuit, with evidence spanning multiple methodological approaches. Early findings from microdialysis studies suggest increased dopamine levels during feeding (4,5); however, the technique lacks sufficient temporal resolution to capture rapid fluctuations. Research utilizing pharmacological and genetic manipulations further demonstrated a crucial role for dopamine in feeding behavior (6–8). For example, dopamine deficient mice will not eat and will die unless administered levodopa (9). While considerable data suggest that dopamine facilitates food-seeking, it has also been proposed that diminished dopamine activity can reduce food intake, with some hypothesizing that diminished dopamine activity mediates satiety and disengagement from eating (10).

Midbrain DA is sensitive to regulation via circulating signals that reflect the animal’s current energetic state and convey post-ingestive feedback, including insulin, leptin, ghrelin, and numerous others (10,11). This sensitivity provides a mechanism whereby modulation of dopamine could link behavioral activation or disengagement to changing physiological energy states. Insulin resistance, a common feature of obesity, can disrupt this feedback loop, potentially impairing a putative role for dopamine in stopping eating in response to satiation (12). The effects of these energy signals on dopamine can be complex and difficult to characterize. For example, insulin effects on dopamine vary in a region and time-dependent manner (11,13), often exerting both positive and negative modulation. The extent to which the complex effects of energy signals might be integrated in the dopamine system to signal the transition from hunger to satiety and, in turn, modulate consumption and meal patterning, has not been evaluated *in vivo*.

Prior studies have examined DA in awake behaving animals during feeding, but these studies are typically in structured operant tasks, employ food restriction, and commonly focus on reward-related behavior rather than changes in appetitive drive or satiation (14–16). Indeed, by employing food restriction and limiting the number of pellets an animal can obtain during a single session of such tasks, the explicit goal is to eliminate the confound of satiation. Moreover, chronic food restriction induces profound changes across neurophysiological regulatory systems, including increased insulin sensitivity, decreased circulating leptin, chronic stress activation, and many others (17,18). Such changes can have significant functional consequences, including altering brain reward stimulation (19–21). Carr and colleagues have demonstrated that food restriction alters both the dopamine system and target regions (22,23).

Recent work using calcium imaging has demonstrated that dopamine activity increases during feeding of a palatable food (24), consistent with prior literature. Zhu *et al* (24) further show that administration of a GLP-1 agonist induces pharmacological satiety, concurrently reducing feeding and DA cell activity; experimentally restoring DA cell activity overrides the GLP-1 induced satiety and restores feeding. These novel findings demonstrate that dopamine activity can oppose satiety signals and sustain hedonic feeding. However, the accompanying reduction in dopamine activity during GLP-1 induced satiety raises the question of whether decreased dopamine signaling normally mediates satiety, as this was not directly tested. In contrast, other findings suggest that GLP-1 administration does not reduce DA (25), leaving the question of how GLP-1 might alter dopamine unresolved.

In short, although it has been robustly demonstrated for decades that dopamine can drive appetitive behavior, whether or not decreased dopamine activity can serve as an effector mechanism for satiety remains an open question. No study has tracked changes in DA release in awake, behaving animals that correspond to progressive satiety in naturalistic feeding, i.e., without food restriction or other manipulations, to determine if dopamine signaling changes with naturally occurring hunger and satiety.

Here, we use fiber photometry in a naturalistic feeding paradigm to track real-time DA across feeding, providing a view of how these signals change with progressive satiation. We then examine how overnight fasting and pharmacological satiation with a glucagon-like peptide-1 (GLP-1) agonist alters DA signaling across feeding. DA activity increases during food approach, as expected. However, DA transients on approach vary within and between meals, primarily in the nucleus accumbens core (NAcc), reflecting short-term fluctuations in hunger and satiety with progressive ingestion. Fasting revealed a dissociation between initiating and terminating meals, with the DA transients we observe and measure during feeding modulating termination of meals. In contrast, pharmacological satiety induced by a GLP-1 agonist did not reduce the magnitude of feeding related DA transients, but blunted variability in DA transients associated with short-term fluctuations in hunger and satiety effectively decoupling DA from meal patterning.

## Materials and Methods

### Animals

Mice of both sexes, approximately 120 days of age were used for all experiments. We originally intended to simultaneously record calcium signals from striatal projection neurons in both pathways. After the initial experiment, we discontinued the GCaMP recording (i.e., in the subsequent fasting and pharmacological satiety studies) and focused on DA in the nucleus accumbens. Thus, in the first experiment, we used selective cre-recombinase lines to target GCaMP to indirect pathway spiny projection neurons (iSPNs; Adora-cre: B6.FVB(Cg)-Tg(Adora2a-Cre)KG139Gsat/Mmucd; GENSAT, 036158-UCD) or direct pathway SPNs (dSPNs; D1-cre: B6.FVB(Cg)-Tg(Drd1a-cre)EY262Gsat/Mmucd; GENSAT, 030989-UCD). In the subsequent experiments testing the effects of fasting and a GLP-1 agonist without GCaMP recording, we used wild-type C57BL/6J mice (Jackson Laboratories, #000664). Mice were singly housed after surgery to protect cranial implants. Mice were maintained on a 12:12 hour light-dark cycle and provided *ad libitum* water and food (either freely available on cage floor before testing or freely available via automated feeder during testing). All procedures were approved by the Queens College Institutional Animal Care and Use Committee.

We report the data and analysis from the GCaMP Ca++ recordings from the initial experiment in supplementary materials (supplemental materials, GCaMP recordings), along with our rationale for separating examination of DA and SPN activities into two separate projects. No differences were observed in dopamine signaling between the Adora-cre and D1-cre lines used in the first experiment (z ΔF/F DA signal pellet at approach, main effect of line, *F*(_1,11_) = 0.017, *p* = 0.89) and so the DA data from these two lines are combined in the analysis reported below.

### Automated feeding paradigm

Mice were habituated to the experimental room and grain pellets for five days prior to testing. Mice were then moved to chambers with automated feeders (FED3, Matikainen-Ankney et al., 2020) that dispensed a single 20 mg grain pellet at a time (Bioserv, cat# F0163). Mice could retrieve pellets *ad libitum* with no work requirement; after each pellet is retrieved, another is immediately made available. Every pellet retrieval was time-stamped and recorded. Mice were allowed 1-2 days to adjust to the automated feeder, and then feeding was monitored for 4 days to assess meal patterns. We recorded with fiber photometry the first four hours after the onset of the active/dark cycle of the 4th day. For fasting experiments, the pellet dispenser was turned off, and no food pellets were available for the fasted group on day three of baseline, prior to recording. The following day, at the onset of the active cycle, the pellet dispensers were turned on, and fiber photometry was recorded.

### Pharmacological satiety

Exendin-4 (Ex-4), a GLP-1 agonist, was used to induce pharmacological satiety (27). Ex-4 (Cat# E7144, Sigma-Aldrich, Saint Louis, MO) was dissolved in 0.9% saline and given either 90 μg/kg (pharmacological satiety group) or saline controls (Veh) via subcutaneous injections 45 minutes prior to onset of the dark cycle. To reduce stress of injections, mice were handled and given ’mock’ injections (touch tip of capped syringe to domed skin) for 3 days prior to Ex-4 administration. In this experiment, we recorded only the first two hours of feeding. Based on previous findings, we expected the lead time to be forty-five minutes post s.c. injection and the effects of Ex-4 on hunger and satiety to last about two hours (27).

### Fiber photometry

In experiment 1, we co-expressed the red dopamine sensor RdLight1 (AAV9-Syn-RdLight1; Cat# 1646-AAV9) and the green calcium sensor GCaMP6f (AAV5.Syn.Flex.GCaMP6f.WPRE.SV40; Cat# 100833-AAV5), targeting GCaMP to either d- or i- SPNs using the D1-cre or Adora-cre mouse lines noted above. 350 nL of the RdLight1/GCaMP6f virus mixture were injected into either the nucleus accumbens core (NAcc; AP: +1.3mm, LAT: +.9mm, DV: −4.25) or shell (NAcSH; AP: +1.3mm, LAT: +.7mm, DV: −4.4) and mice were allowed three weeks for recovery and expression of the sensors. Fluorescent signals were recorded using a Neurophotometrics dual-color fiber photometry system.

For signal processing, RdLight1, GCaMP6f, and isosbestic signals were de-interleaved. We denoise each signal by computing the simple moving average with a 10-data-point window. We correct for photobleaching with a moving mean algorithm using reweighted penalized least squares (28,29). Signals are standardized using Z-scores. To calculate ΔF/F signals and correct for movement artifacts, isosbestic control signals were fitted, scaled, and subtracted from RdLight1 and GCaMP6f signals, then divided by the fitted isosbestic signal (29). Due to the extended 4-hour recording period, we adjusted the LED’s excitation power to 40 µW, reduced the sampling rate to 20 Hz, and incorporated blank frames into the duty cycle to enhance our signal-to-noise ratio and minimize photobleaching. We used only fiber optic implants with a greater than 85% light efficiency (30,31).

### Statistical Analyses

Signal processing was performed using a custom-written script in MATLAB (RRID:SCR_001622, Mathworks Inc., MA). All other analyses, statistics, and graphs were generated using R Project for Statistical Computing, v. 4.4.1 (RRID:SCR_001905).

ANOVA was used to test the significance of the behavior data, followed by a Tukey post-hoc pairwise comparison. The statistical significance of fiber photometry data was evaluated using linear mixed-effects models (LMM) with the lme4 R package (RRID:SCR_015654, (32). The LMM included individual mice as a random variable. *p* values and degrees of freedom were calculated from LMM models using Satterthwaite’s degrees of freedom method in the lmerTest

R package. We used a mixed logistic regression (MLR, estimated using maximum likelihood and the Nelder-Mead optimizer) to predict binary outcomes (i.e., whether the mouse takes a next pellet), again with mouse as random variable. 95% Confidence Intervals (CIs) and p-values were computed using a Wald z-distribution approximation. Although we used both male and female mice, the study was underpowered to effectively evaluate sex differences. We combined sexes in all analyses below and perform a targeted but statistically limited analysis of sex differences separately, available in supplemental material.

## Results

### Dopamine activity increases during food approach

We used fiber photometry in mice to measure real-time changes in extracellular DA in the striatum across feeding phases, including approach, consumption, and post-consumption (recovery) in an automated free-feeding paradigm (Figure 1A). Mice could collect a 20 mg pellet at will, after which a new pellet was immediately dispensed and available. The phases of feeding were defined as windows of time relative to pellet retrieval (time 0). The baseline was defined as -10 to -2 s before pellet retrieval; food approach as the 2s immediately prior to pellet retrieval (-2 to 0), consumption was 1 to 3 s post pellet retrieval, and recovery was 6 to 10 s post pellet retrieval, similar to (31).

**Figure 1.**
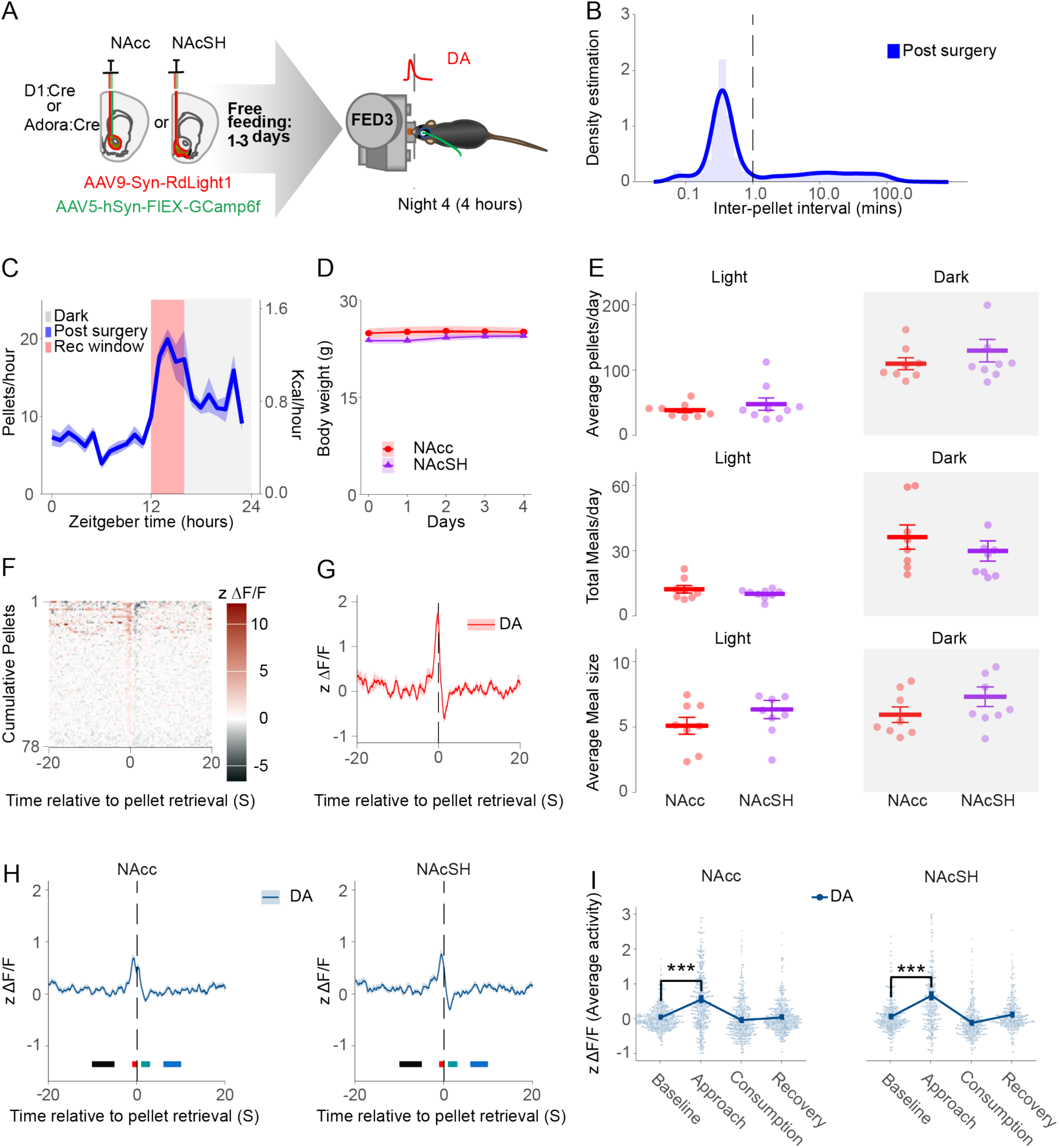
DA activity increases during food approach in naturalistic feeding. **A)** Schematic of recordings of DA activity for 4 hours at the onset of the dark cycle in free-feeding mice. **B)** Interpellet interval density plots for free-feeding (defining a meal). **C)** Mean number of pellets/hr across a 24-hr period (all mice). **D)** Body weights over four days by groups. **E)** The total number of pellets, meals, and average meal size for all four days of feeding by groups and cycle. **F**) Heatmap of DA release (Z scores) relative to pellet retrieval across cumulative pellets for an example mouse. **G**) Mean DA release relative to pellet from the example mouse in 1F **H)** Average dopamine release (Z scores) relative to pellet retrieval by sub-region of the nucleus accumbens. Black, red, green, and blue rectangles indicate feeding stages: baseline, approach, consumption, and recovery, respectively. **I)** Peak DA release during feeding stages and sub-regions. Data are presented as the mean ± s.e.m. in \**p* < 0.05; \*\**p* < 0.01; \*\*\**p* < 0.001.

Examining the interpellet interval across all mice (Figure 1B), most pellets were consumed within a minute. We defined a ’meal’ as a string of consecutive pellet retrievals with interpellet intervals of less than one minute. A break longer than one minute from retrieval was considered the start of a new meal (33). Consistent with prior studies (33), food intake at the onset of the dark cycle peaked at approximately 20 pellets/hour, during which mice consume, on average, 44.76 ± 3.5 pellets over the first 4 hours of the night cycle. There was little variation across mice in pellets eaten during 24-hour periods. We recorded this first 4 hours (shaded red rectangle, Figure 1C).

We compared mouse groups distinguished by subregion targeted for photometry, i.e., either nucleus accumbens core (NAcc) or the nucleus accumbens shell (NAcSH). Neither body weight (Fig 1D) or eating patterns (Figure 1E), including total consumption (pellets per day), the number of meals per day and average meal size differed across these groups. Mice exhibited stable feeding patterns across all groups and days.

DA release increases during pellet approach, i.e., immediately preceding pellet retrieval (Fig 1F-G shows an example mouse). This increase was consistent across mice and observed in both the NAcc and NAcSH with no significant differences between the regions and no changes observed during the other phases of feeding (Figure 1H-I, blue traces, approach vs. baseline, *F*_(3, 2706)_ = 126.6, *p* < .001). These data indicate that transient increases in DA activity correlate with food approach rather than consumption.

### Dopamine activity attenuates as mice reach satiation in the NAcc

We inspected the quality of the DA traces for artifacts and an appropriate signal-to-noise ratio; most transient peaks exceeded one z-score (blue triangles in example trace, Figure 2A). We observed progressive changes in DA release occurring during food approach within individual meals. Specifically, DA release declines within a meal (Figure 2A, red traces) across consecutive food approaches (Figure 2A, bottom panel, black triangles). As meal lengths varied, we summarized changes in dopamine signals at each pellet retrieval across consecutive pellets within a meal, separating meals of different sizes (Figure 2B). In the NAcc, we observe a general pattern of peak DA release at food approach, which declines across individual meals (red trace), an effect most prominent in both shorter and longer meals. In contrast, in the NAcSH (purple trace), DA release at food approach did not exhibit a consistent pattern across pellets within individual meals. We quantified these changes in peak DA by comparing the first and last pellets of meals. Consistent with the patterns observed in Figure 2A, we found a significant difference in peak DA release at food approach from the first to the last pellet of a meal (Fig 2C, interaction between pellet position x phase (approach) x region, *F*(_3,1035_) = 3.01, *p* = 0.029), with post-hoc showing this reached significance only in the NAcc (Tukey, *t*(_1035_) = 5.02, *p* < .001). These data demonstrate a short-term, progressive decrease in peak DA during pellet approach from initiation to termination of a meal, consistent with short-term satiety; that is, progressive ingestion of pellets attenuates DA signaling during food approach, suggesting declining appetitive motivation.

**Figure 2.**
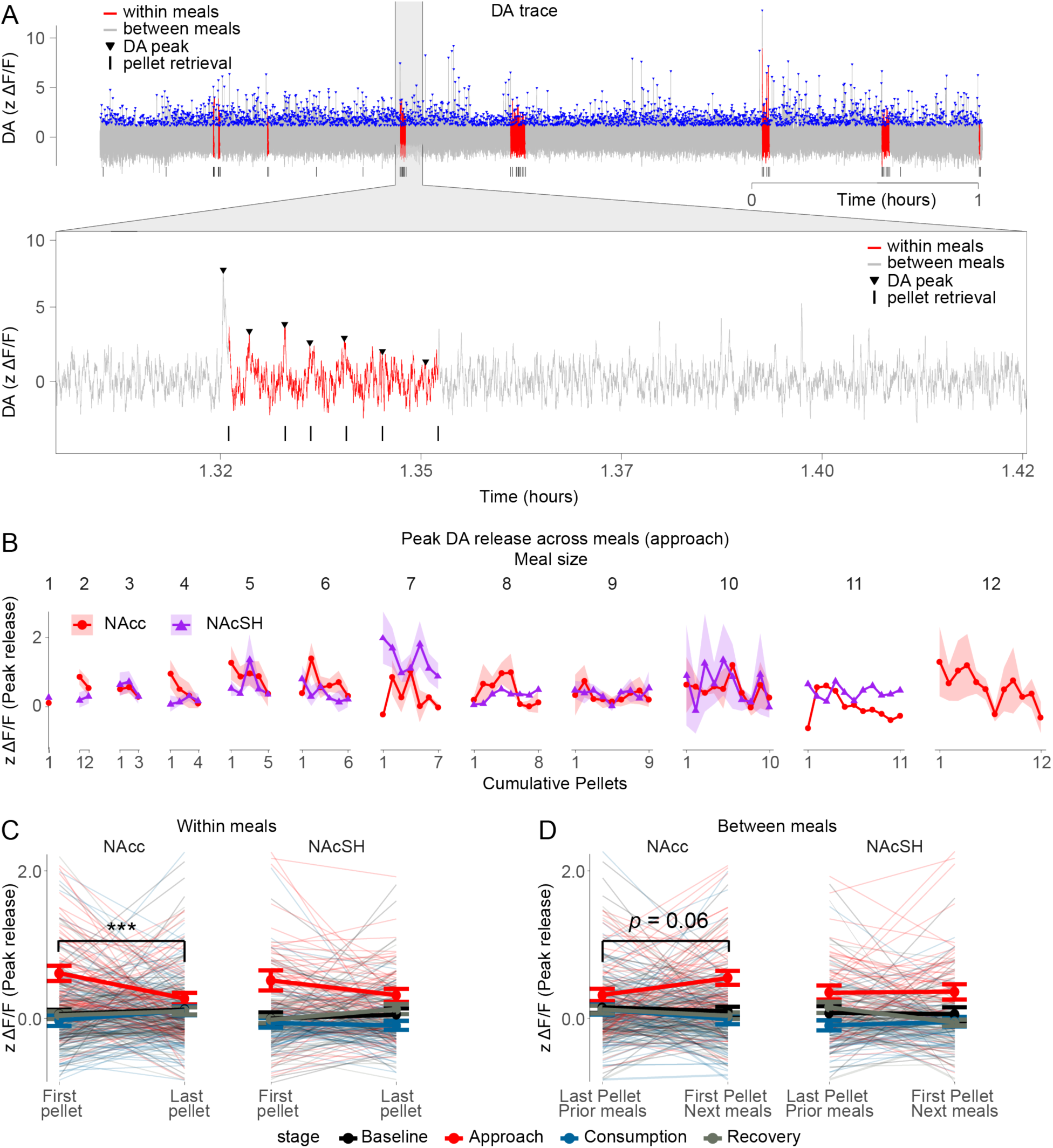
Dopamine activity attenuated as mice reached satiation in the NAcc. **A)** Top: An example DA trace across time (top), red and grey, indicating that DA occurred within or between meals, respectively. Blue down-facing triangles indicated DA peaks above the threshold of 1. Bottom: A zoomed-in plot showing the progressive decline of DA release; the vertical black line indicates pellet retrieval. **B**) The average DA release in the NAcc and NAcSH was grouped by meal size across cumulative pellets during food approach. **C**) Average DA activity from the first and last pellet of a meal, grouped by the NAcc and NAcSh. **D**) Average DA release from the last (prior meal) and first pellet (next meal) between meals grouped by the NAcc and NAcSH.

To determine if the decline in DA between meals recovered, we compared peak DA at pellet approach between the last pellet of a meal to the first pellet of the subsequent meal. Peak DA release at the first pellet of a meal was increased compared to the last pellet of the previous meal, suggesting that the strength of the DA signal on approach recovered between meals, (Figure 2D, interaction between pellet position x event (approach) x region, *F*_(3,915)_ = 3.32, *p* = 0.019), though post-hoc comparing regions indicates marginally significance only in the NAcc (Tukey, *t*_(915)_ = 3.34, *p* = 0.066). This recovery of peak DA at food approach on initiating a new meal suggests an underlying diminution of short-term satiety.

We examined whether the magnitude of peak DA release on a meal’s first pellet corresponds to the meal’s length, i.e., whether a higher initial peak leads to longer meals. We first compared this DA peak from all meals larger than one pellet to single pellet meals. Peak DA at approach is significantly higher in multi-pellet compared to single-pellet meals in the NAcc (Figure 3A, Tukey: *t*_(220)_ = 3.98, *p* < 0.01) but not the NAcSH (Figure 3A, Tukey: *t*_(216)_ = 1.37, *p* = 0.518). This suggests that the magnitude of DA release at the beginning of a meal could influence meal length.

**Figure 3.**
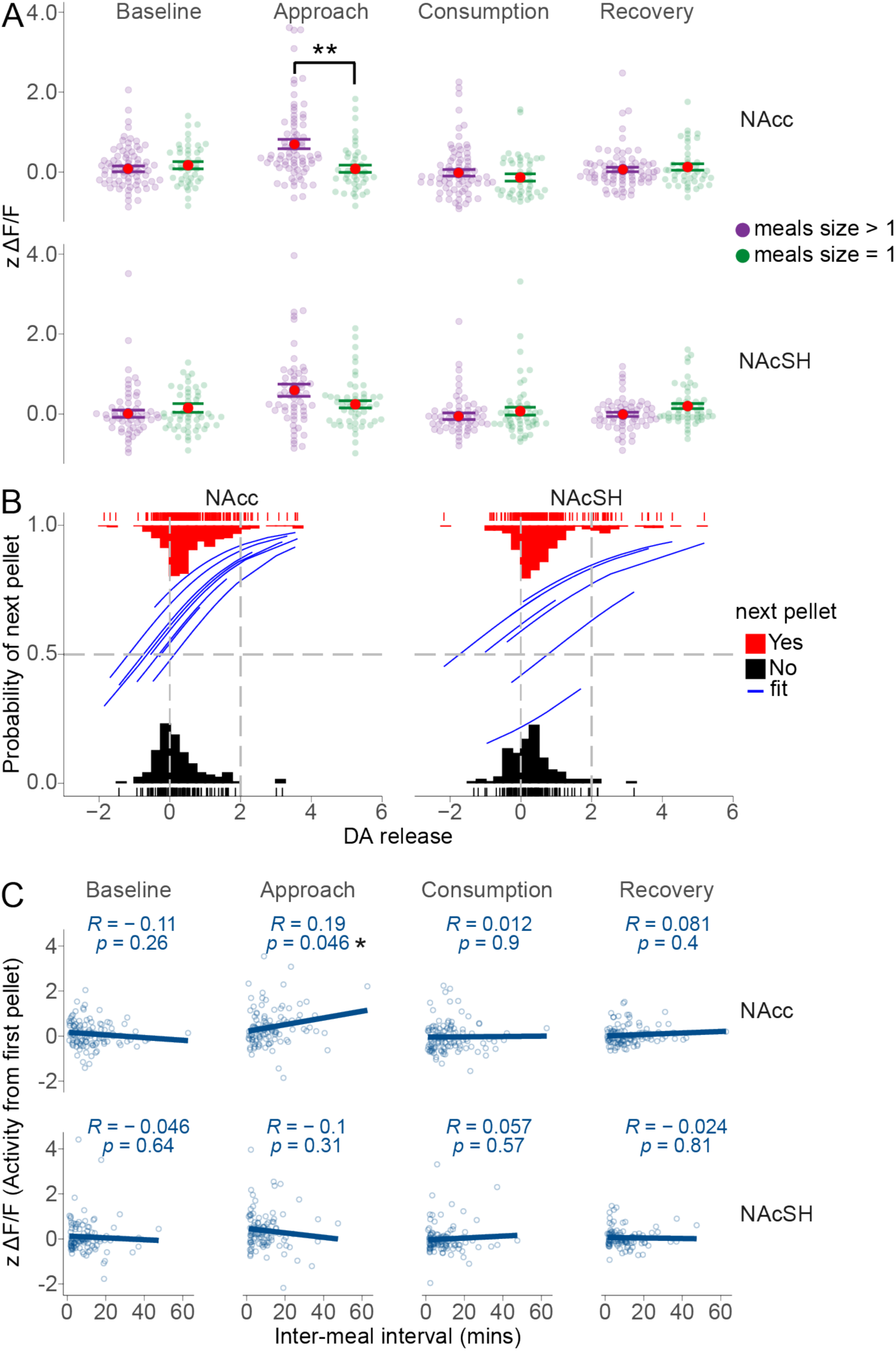
Dopamine release influences meal patterns**. A)** The DA release of meals larger than one pellet (Mult) compared to meals that are exactly one pellet (single) for baseline, approach, consumption, and recovery. **B)** A mixed logistic regression model showing that peak DA release significantly predicts the probability of the next food pellet in the NAcc. Blue traces represent the fit of each mouse. **C)** Linear regression of the average DA release for the first pellet of a meal during food approach predicted by the inter-meal interval. Error bars: ± s.e.m.; \**p* < 0.05; \*\**p* < 0.01; \*\*\**p* < 0.001.

We tested whether the initial DA peak correlates with meal length for each feeding stage. Unexpectedly, it does not (*R* = -0.039, *p* = 0.658). To further probe the relationship between DA release and meal size/length, we tested whether DA release facilitates subsequent pellets, leading to longer meals. To achieve this, we estimated the relationship between DA and the binary outcome of taking or not taking a next pellet, accounting for the within-subject nature of our data using a mixed logistic regression. Higher DA release on approach to pellet retrieval significantly increased the probability of a next food pellet being taken (blue traces indicate each mouse predicted fit) in the same meal (Figure 3B; OR = 1.84, 95% CI: 1.4-2.42, *p* < 0.001) with no difference between regions.

We examined whether the initial DA peak at the beginning of a meal was influenced by the length of time since the last meal. Consistent with the recovery of DA peak at the first pellet of a new meal (i.e., Figure 2D), DA peaks increase with time since termination of the previous meal in the NAcc (Figure 3C top, *F*_(1, 108)_ = 4.09, *p* < 0.05) but not in the shell (Figure 3C bottom, *F*_(1, 101)_ = 1.06, *p* = 0.306). No relationship was observed between DA signals and the length of time since the last meal in other segments of the signal (i.e., baseline, consumption, or recovery).

These data demonstrate a relationship between the magnitude of DA transients associated with food approach and the regulation of eating on a minute-to-minute timescale. Within a meal, peak dopamine declines with successive pellet consumption, consistent with short-term satiety. The longer the mouse waits to eat, the higher the DA release is at the start of a meal. Moreover, greater DA release facilitates taking a next pellet, extending the size of the meal, though this effect is probabilistic. Thus, overall, the higher the DA release, the more likely the mouse is to continue eating. We next tested the effects of a 24-hour fast and pharmacological satiety (GLP-1 agonist) on these signals and their relationship to meal patterns.

Across the defined phases (i.e., baseline, approach, consumption and recovery), the most robust effects were evident in approach, with neither consumption or recovery differing from baseline (Fig 1I, 2C-D, 3A,C). In remaining analyses, we included either approach only or approach plus baseline for comparison, depending on the specific analysis.

### Hunger modulates DA activity associated with food approach

In this and the GLP-1 studies below, we examined only the NAcc. To assess how hunger modulates eating behavior and underlying striatal dopamine signaling, we fasted mice for 24 hours (Figure 4A), resulting in a short-term, modest loss of body weight that recovered the following day (Figure 4B). We compared the non-fasted controls from this experiment to the mice in the first experiment where we recorded from the NAcc (Figure 2, reported above) and observed no significant differences in their meal patterns (data not shown). We found no significant difference in DA signaling at pellet approach (main effect of experiment, *F*_(1,9)_ = 0.07, *p* = .791) and therefore combined the non-fasted control data with the NAcc data from the prior experiment for comparison with fasted mice.

**Figure 4.**
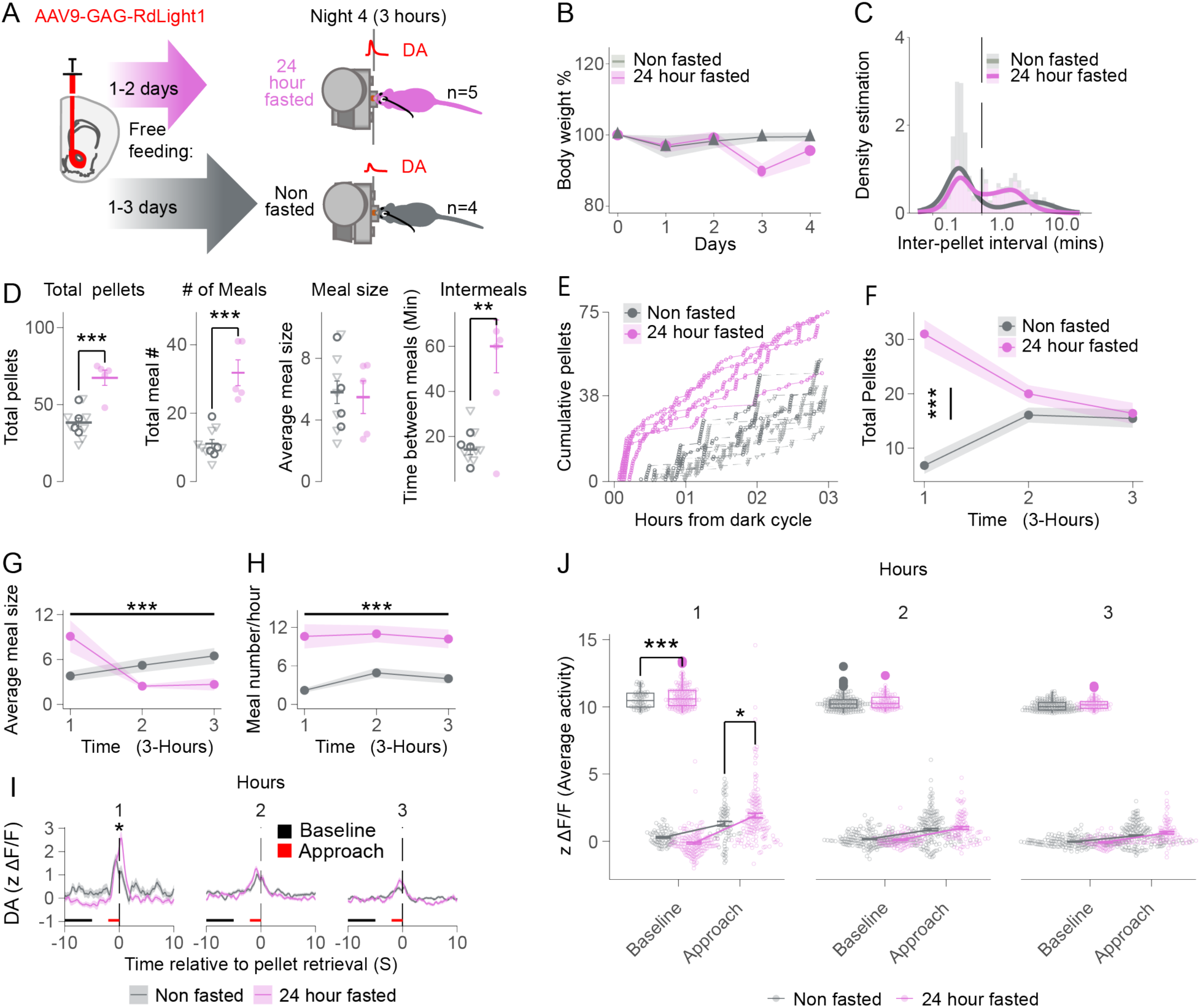
A 24-hour fast increases DA release during the first hour of feeding. **A)** (left) Surgery for expressing RdLight1 in WTB6 mice in the NAcc, (right) Meal patterns assessed using FED3 for 24 hours across two days, then half the mice fasted 24 hours, and the other half were left on free feeding. On night 4, at the onset of the dark cycle, DA release was measured in both fast and 24-hour fast mice for 3 hours. **B)** Bodyweight percent by groups across days. **C)** Interpellet interval density during recordings (2 hours). **D)** The total number of pellets, meals, average meal size, and intermeal time for 3 hours of recordings. **E)** Cumulative pellets eaten across 3 days for each mouse, grouped by whether they were fasted. **F)** Total pellet intake by the hour for each group. **G)** Mean meal size across hours by group. **H)** Meal frequency across hours by group. **I)** The average DA release in Z scores, grouped by whether the mice were fasted, by hour. **J)** Average activity of DA release during baseline and food approach for both groups by the hour. The inset is the slope of the baseline and approach. Error bars: ± s.e.m.; \**p* < 0.05; \*\**p* < 0.01; \*\*\**p* < 0.001. 24-hour fast, n=5, Not fasted, n=10. Triangle markers indicate previous controls.

The patterning of meals, i.e., interpellet interval distribution, was similar between the fasted and non-fasted mice and aligned with the previous experiment (Figure 4C). However, the small peak at a longer interval, presumably reflecting the most common intermeal interval, is slightly left-shifted and higher in fasted mice, suggesting a shorter period between meals and an increase in meal frequency. There was no difference in the average interpellet interval within meals as defined here. Increased hunger led to higher food intake. The 24-hour fasted mice consumed nearly double the pellets in 3 hours compared to the non-fasted mice (Figure 4D, pellets: *F*_(1, 14)_ = 29.75 *p* < .001), eating twice as many meals (Figure 4D, meals: *F*_(1, 14)_ = 46.10, *p* < .001). The average size of individual meals appeared unchanged during the 3-hour period (Figure 4D, meal size: *F*_(1, 14)_ = 0.06, *p* = 0.80). Consistent with a greater frequency of meals, the intermeal interval was significantly decreased in the fasted mice (Figure 4D, intermeals: *F*_(1, 14)_ = 13.80, *p* = .002).

We evaluated whether fasted and non-fasted mice exhibited different meal patterns over successive hours, reflecting accumulated consumption. The increased consumption in the fasted mice occurred almost entirely within the first hour (Figure 4E-F, Tukey: *t*_(42)_ = -8.99, *p* < 0.001). Although meal size averaged across 3 hours is unchanged between groups, meal size substantially varied by hour in the fasted compared to non-fasted mice (Figure 4G, group x hour: *F*_(2, 28)_ = 14.13, *p* < .001). Specifically, the fasted mice consumed larger meals in the first hour and then smaller meals in the second two hours. Despite these changes in meal size by hour, the number of meals taken remained consistently elevated across all three hours in the fasted compared to non-fasted mice (Figure 4H, group, *F*_(1, 43)_ = 20.04, *p* < .001). Thus, in the first hour, both meal size and meal number are contributing to greatly increased consumption, while in hours 2-3 meal number remains elevated while meal size decreases to less than that seen in non-fasted mice, resulting in comparable consumption between the groups in those hours (Figure 4F, hr 2, Tukey: *t*_(42)_ = -1.45, *p* = 0.69; hr 3, Tukey: *t*_(42)_ = 0.352, *p* = .99).

These data suggest that fasting induced an elevated energy deficit at the beginning of the active cycle that drove compensatory consumption and altered meal patterning, i.e., the starting and stopping of meals. This observed compensatory eating comprises two dissociable components. First, we observe an increase in meal frequency across the entire 3-hour period recorded, suggesting that the energy deficit generated by overnight fasting leads to more rapid recovery of hunger after short-term satiation resulting in an increased tendency to initiate a new meal; that is, the mice *start* eating again sooner. Second, we observe changes in meal size that vary hour-by-hour. In the initial hour, meal size is substantially larger, reflecting delayed termination of meals. This, in combination with more frequent meals, mediates substantial compensatory eating. In hours 2-3, however, meal size decreases substantially, reflecting an *earlier* termination of eating. This earlier termination of meals resulted in smaller meals than non-fasted mice. The impression is of two separate processes: one mediating ’interest’ in eating (meal initiation), which appears to be elevated in a sustained way and the other mediating feedback on the necessity of continued eating, i.e., satiety. In the second and third hours, elevated ’interest’ and greater frequency of meal initiation is apparently offset by more rapid satiety resulting in a ’check’ on fasting induced compensatory (over-)eating.

DA release differed between fasted and non-fasted mice only on approach during the first hour (Figures 4I-J, hour x group x baseline vs. approach, *F*_(2, 1492)_ = 5.91, *p* < .01). We quantified differences between the fasted and non-fasted mice by comparing the slopes of DA increases from baseline to approach across hours 1 through 3. The 24 hour fasted mice exhibit significantly greater slopes compared to non-fasted mice in the first hour (Figure 4J, top inset, Tukey: *t*_(753)_ = -4.89, *p* < .001) but not in hour 2 or 3 (Figure 4J, top insets, hr 2:Tukey: *t*_(753)_ = -.92, *p* = 0.94; hr 3: *t*_(753)_ = -1.283, *p* = 0.79). This enhanced DA peak at approach in fasted mice during the first hour only is consistent with the increased consumption in the first hour and, moreover, suggest that this increased dopamine specifically corresponds to prolonged meals, or conversely diminished satiety and meal termination. In contrast, increased frequency of meal initiation occurs in hours 2 and 3 in the absence of any detectable difference in dopamine between the fasted and non-fasted mice in the NAcc.

We evaluated whether a 24-hour fast impacts the progressive attenuation of DA as individual pellets are consumed within a meal. Comparing DA release between the first and last pellet of a meal, we observe as before, a decrease in peak dopamine on pellet approach for both fasted (Figure 5A, Tukey: *t*_(578)_ = 4.93, *p* < .001) and non-fasted mice (Figure 5A, Tukey: *t*_(578)_ = 4.09, *p* < 0.01). Similarly, as above, both the fasted mice (Figure 5B, right, Tukey: *t*_(514)_ = 4.01, *p* < .01) and non-fasted mice (Figure 5B, right, Tukey: *t*_(514)_ = 3.53, *p* < .05) showed significant recovery of DA peak on pellet approach-retrieval between meals.

**Figure 5.**
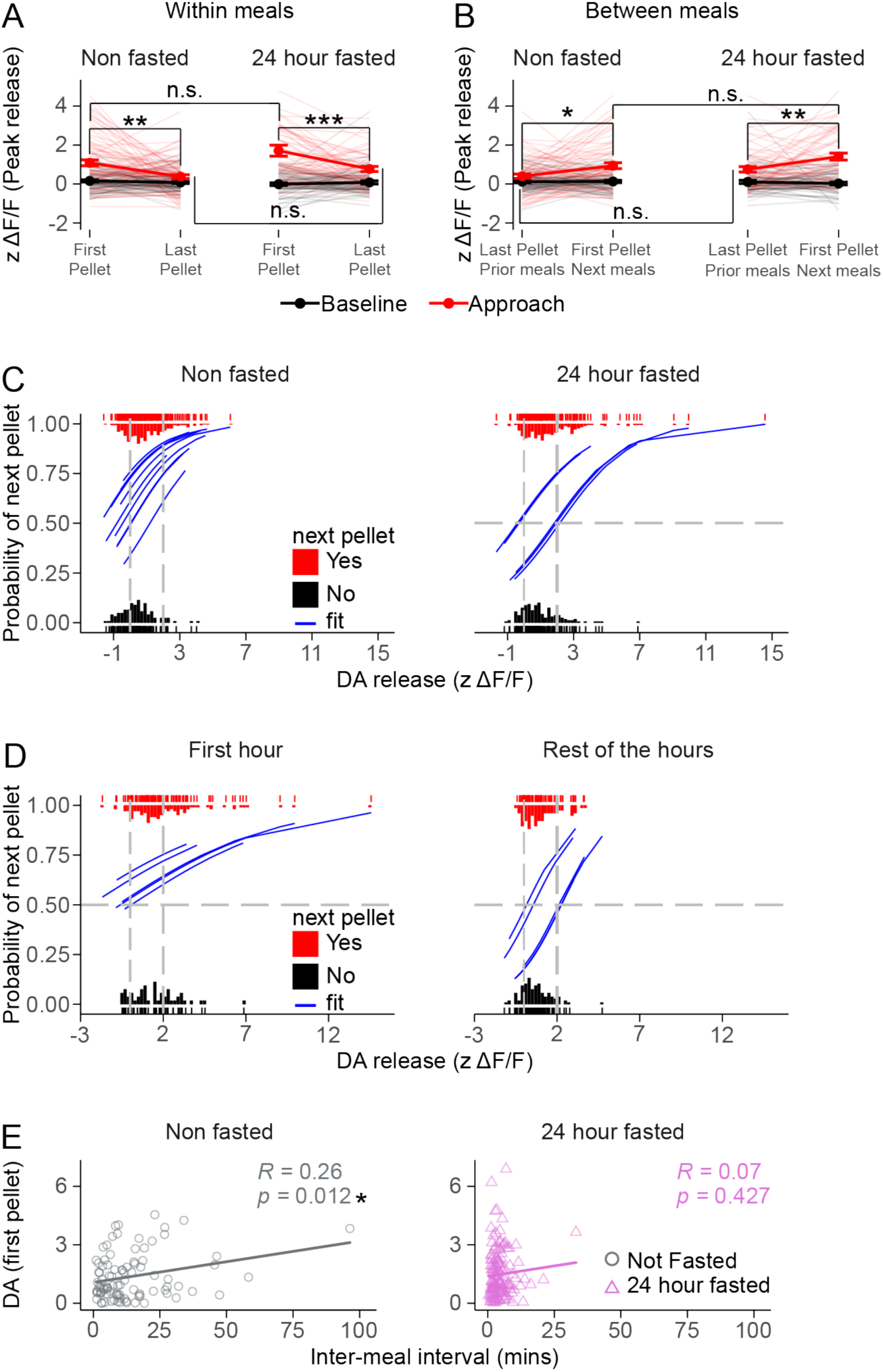
A 24-hour fast reduces dopamine’s influence on meal size. **A)** Average DA activity from the first and last pellet of a meal, grouped by the fasted condition. **B)** Average DA release from the last and first pellet between meals grouped by the fasted condition. **C)** A mixed logistic regression showing peak DA release significantly predicts the probability of the next food pellet in non-fasted and fasted mice. Blue traces represent the fit of each mouse. **D)** In the fasted condition, A mixed logistic regression analysis showed that the peak DA release probability is shifted according to the time of food consumption. **E)** DA release during the first pellet of a meal regressed against time since last meal. Blue traces represent the fit for each mouse, grouped by fasted condition. Error bars: ± s.e.m. in \**p* < 0.05; \*\**p* < 0.01; \*\*\**p* < 0.001. 24-hour fast, n=5, Not fasted, n=10.

Surprisingly, we did not observe a difference between fasted and non-fasted mice in this within meal decline in DA peaks between the first and last pellet of a meal, nor with recovery after an intermeal interval. However, as the number of pellets in a meal differs between fasted and non-fasted (Fig 4G, greater in hour 1, less in hour 2-3), this suggests that the rate of decline in DA transients on approach from one pellet to the next with a meal must also differ, and similarly as intermeal intervals differ between groups, that the rate of recovery must also differ.

We examined whether the magnitude of the dopamine peak at approach mediates the likelihood of taking a next pellet. When we examine this for the entire period, greater DA release at approach increased the probability of the mouse taking a next pellet for both fasted and non-fasted mice (Figure 5C, z ΔF/F, OR = 1.73, 95% CI: 1.34 - 2.23, *p* < 0.001; z ΔF/F x group, OR = .92, CI: 0.l67-1.27, *p* = .62). There was a modest main effect of fasting independent of dopamine (OR = .36, 95% CI: 0.16 - 0.78, *p* = 0.01), not surprisingly indicating a non-dopamine mediated process that modulates likelihood of taking a next pellet, though surprisingly small compared to the effect of peak DA. Because the fasted mice exhibit different eating patterns across hours, we repeated this analysis with the fasted mice only separating out the first from the remaining hours (Fig 5D). In these fasted mice, during the first hour all magnitudes of DA transients during approach were associated with a greater than 50% chance of a next pellet whereas in hours 2-3 the likelihood of a next pellet was decreased across most DA peak magnitudes (Figure 5D, z ΔF/F x hours, OR: 1.45, CI: 0.96 - 2.17, *p* = .07). As above, there is a modest effect of fasting independent of DA (hours main effect, OR: 0.3, CI: 0.16 - 0.59, *p* < .001). Neither hours nor DA x hours were significant in non-fasted mice, only DA predicted a next pellet (OR: 1.71, CI: 1.12 - 2.62, *p* = .01). These results are consistent with larger meals in the first hour and smaller meals in hours 2-3 compared to non-fasted mice.

Finally, we examined the relationship between the amount of time elapsed from the end of the previous meal and the magnitude of the dopamine transient on approach to the first pellet of the next meal (Figure 5E). As above, we observe a linear relationship in the non-fasted mice (Figure 5E, left, *F*_(1, 120)_ = 5.07, *p* < .05), such that the longer the time elapsed since the last pellet, the higher the dopamine peak on the first pellet when the mouse resumes eating. This relationship is lost in fasted mice (Figure 5E, right, *F*_(1, 153)_ = 0, *p* = 0.99).

Overnight fasting altered feeding behavior primarily by increasing meal frequency across the entire recording period and transiently increasing meal size early in the session, reflecting an elevated drive to initiate eating and a delayed termination of meals during the first hour, but earlier termination in hours 2-3. Differences in dopamine signals were observed only during hour 1 when both meal size and frequency were increased, but not during hours 2-3 when meal frequency remained elevated but meal size was reduced compared to non-fasted mice.

Two aspects of dopamine signaling need to be considered, its underlying regulation and its relation to behavior. First, because the pattern of within meal decline in dopamine transients is similar in fasted mice, but the number of pellets in a meal increased (or decreased in hours 2-3), this suggest that fasting has altered the *rate* of decline across these transients within a meal. Similarly, as DA recovery between meals is also similar, but the fasted mice exhibit shorter intermeal intervals, this again suggest that fasting alters the underlying rate of recovery in DA signaling. Together, these observations indicate that fasting alters how short-term changes in hunger and satiety modulate DA signaling.

Second, we observe a specific alteration between DA and behavior. The magnitude of the DA transient on food approach modulates the likelihood of taking a next pellet. In fasted mice, this relationship is enhanced in hour 1, with DA increasing the likelihood of a next pellet and generating larger meals. In hours 2-3, in contrast, this relationship is diminished, with DA less likely to predict a next pellet, consistent with shorter, earlier termination of meals observed. This latter effect likely arises not from fasting per se, but as a result of the compensatory eating that occurred during hour 1.

### Pharmacological satiety blunts DA fluctuations within and between meals

In the study above, we ask how increased hunger alters the relationship between dopamine signaling, eating and meal patterning. Here we ask the opposite, how does decreased hunger alter these relationships? A traditional approach would be to pre-feed the animals; however, doing so in a naturalistic context (not session-based operant) leads to animals who subsequently eat little to nothing, providing insufficient analyzable data until their hunger returns, eliminating the sated state that was the aim of pre-feeding. This, in essence, is simply our experiment in reverse: studying changes in DA signaling over the course of hunger *returning* rather than progressive satiety; however, unlike satiety, which we can mark by successive pellet consumption, in returning hunger we have no visible marker of when animals get incrementally more hungry. Instead, we opted for pharmacologically induced satiety administering a GLP-1 agonist Exendin-4 (Ex-4) that demonstrably reduces consumption (27). Moreover, the mechanisms by which GLP-1 modulates eating, including the modulation of dopamine (24), are of importance given the dramatic increase in the use of these agents as pharmaceuticals. Here we ask how GLP-1 induced alterations in naturalistic eating might correspond to changes in dopamine signaling. In contrast to pre-fed animals, these animals do eat and provide data for analysis, but they eat less. Our goal is to assess the potential role of altered dopamine in this reduced consumption. For example, does GLP-1 accelerate the progressively decreasing dopamine transients we observed above with successive pellet approaches, thus shortening meals and reducing overall consumption?

Based on prior literature (27), we administered Ex-4 45 minutes prior to the onset of the dark cycle to achieve 1.5-2 hours of pharmacological satiety during the initial onset of feeding (Figure 6A). Ex-4 administration did not significantly change the distribution of the interpellet intervals (Figure 6B). The initial rate of consumption remained unchanged between Ex-4 and Veh, with the effect of Ex-4 observable in the second hour from the onset of the dark cycle (Figure 6C-D, drug x hour, F_(1,14)_ = 12.81, *p* < .01). Ex-4 administration robustly decreased total pellet consumption and meal size (Figure 6E, total pellets, *F*_(1, 10)_ = 15.28, *p* < .01; meal size, *F*_(1, 9)_ = 7.2, *p* < .05). The observed decrease in the number of meals was marginally significant (*F*_(1,10)_ = 3.99, *p* = 0.07). The apparent increase in intermeal intervals did not reach statistical significance (Figure 6E, *F*_(1, 10)_ = 1.53, *p* = 0.24).

**Figure 6.**
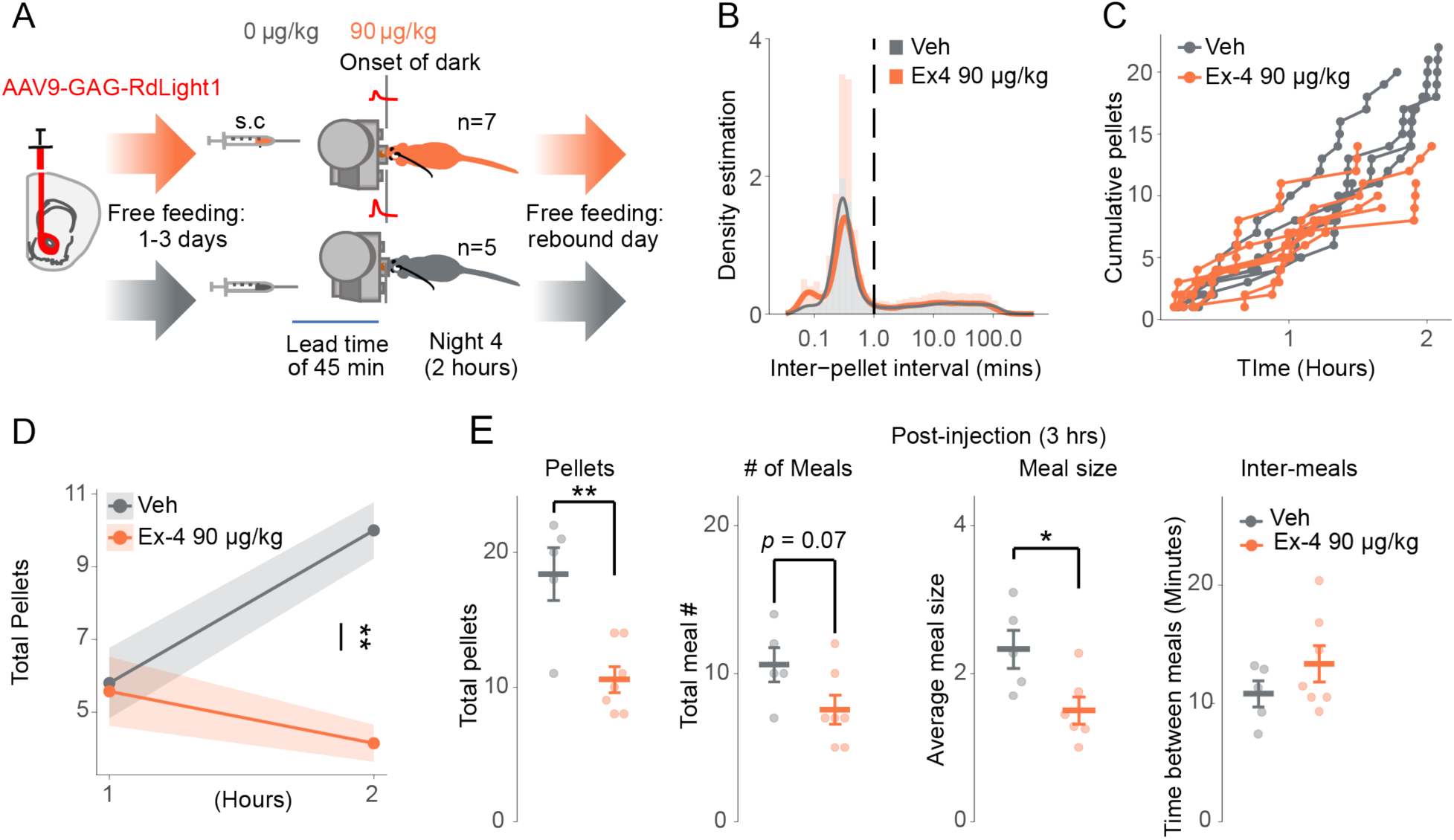
Ex-4 decreases natural food consumption and meal size. **A)** (left), Surgery for expressing RdLight1 in WTB6 mice in the NAcc, (right) Meal patterns assessed using FED3 for 24 hours across 3 days. On night 4, Ex-4 or Veh was injected 45 minutes before the onset of the dark cycle, and DA release was measured for a total time of 2 hours. **B)** Inter-pellet interval density during recordings (2 hours). **C)** The cumulative pellets eaten across time for each mouse, with colored lines indicating Veh or Ex-4 at 90 μg/kg. **D)** The average number of pellets, meals, and meal size 3 hours post-injection on night four. **E)** The total number of pellets, meals, average meal size, and inter-meal interval post i.p. injections. Error bars: ± s.e.m.; \**p* < 0.05; ** *p* < 0.01; *** *p* < 0.001.

We observed no difference in the DA release between Ex-4 and Veh during baseline or pellet approach in either the first or second hour (Figure 7A-B). This is inconsistent with the behavioral data that shows a decrease in consumption during the second hour. We examined whether Ex-4 changes DA dynamics from the first pellet to the last pellet of a meal. In Veh mice, DA levels attenuate from the first to the last pellet of a meal (Figure 7C, left; Tukey: *t*_(115)_ = 3.93, *p* < .01), as observed above. In contrast, this attenuation is diminished in mice administered Ex-4 (Figure 7C, left, Tukey: *t*_(115)_ = .647, *p* = 0.99). Also as above, peak DA at pellet approach significantly recovered between meals in Veh mice (Figure 7C, right, Tukey: *t*_(87.9)_ = 3.44, *p* < 0.05) but not Ex-4 mice (Figure 7C, between meals, Tukey: *t*_(87.9)_ = -0.033, *p* = 1.0). These data suggest that although Ex-4 does not alter the average peak DA on pellet approach (Figure 7A-B), it flattens out within and between meal modulation of these peaks (Figure 7C).

**Figure 7.**
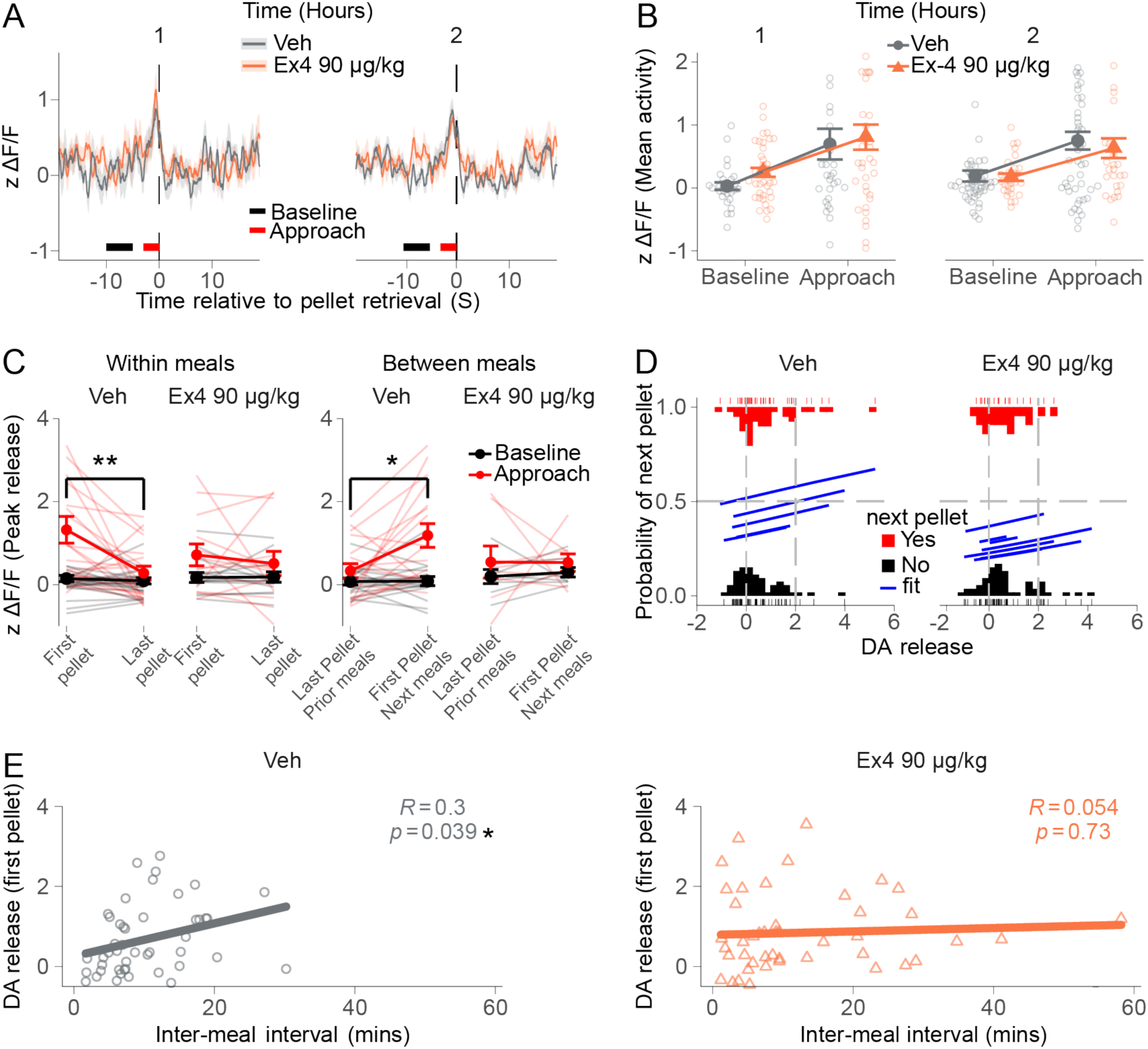
Ex-4 blunts DA attenuation as mice reach satiation. **A)** The average DA release per hour for Veh and Ex-4 90 μg/kg. **B)** The average DA release per hour for each dose, for both the baseline and approach. **C)** The average DA release from the first and last pellets of a meal is grouped by the Veh and Ex-4 at 90 μg/kg, within meals left, between meals right. **D)** A mixed logistic regression model showing the relationship between DA and the probability of the next food pellet in the Ex-4 group. **E)** Linear regression of the average DA release for the first pellet of a meal during food approach, predicted by the inter-meal interval for Veh and Ex-4 at 90 μg/kg. Error bars: ± s.e.m.; \**p* < 0.05; \*\**p* < 0.01; \*\*\**p* < 0.001

We tested whether DA release increases the probability of taking a next pellet. Unlike the studies above, in this experiment DA was not a significant predictor of taking a next pellet. Indeed, the probability of a next pellet was largely below 50%, an effect more pronounced in Ex-4 treated mice (Fig 7D, no main effect of DA, drug, or DA x interaction). Unlike the previous experiments, here all mice were tested under both vehicle and Ex-4 administration and the duration of recording was reduced to two hours. The loss of DA prediction of a next pellet likely arises from a reduced sample number and reduced variability due to using the same mice across conditions.

We performed a linear regression to assess if Ex-4 disturbs the relationship between the elapsed time since the previous meal and the peak DA observed at the first pellet of the next meal. As above, the intermeal interval predicts the magnitude of DA release for the first pellet of a meal for mice administered Veh (Figure 7E, left, *F*_(1, 44)_ = 4.5, *p* < .05). This relationship is lost in mice administered Ex-4 (Figure 7E, right, Ex-4: *F*_(1, 43)_ = 0.12, *p* = 0.72).

In summary, Ex-4 reduced consumption, an effect observed primarily in the second hour, suggesting enhanced satiety. Surprisingly, average DA release associated with pellet approach showed no significant differences between Ex-4 and Veh groups. However, the dynamic regulation of the magnitude of these DA transients within and between meals was substantially diminished. The GLP-1 agonist appears to decouple the modulation of dopamine activity from changing states of hunger and satiety associated with on-going feeding, including short-term satiety during consumption and recovery of hunger between bouts of eating.

Moreover, the relationship between the magnitude of dopamine transient activity and likelihood of continuing to eat is diminished, again suggesting a decoupling of DA from its modulating effects on eating behavior. Thus, similar to fasting, GLP-1 induced satiety acts both on the underlying regulation of dopamine by short-term fluctuations in hunger and satiety, as well in how dopamine modulates feeding behavior. While fasting altered both of these, GLP-1 seems to decouple both from dopamine, suggesting that GLP-1 *disengages* the dopamine system from its normal modulatory role in feeding.

## Discussion

During naturalistic feeding without food restriction or work requirements, we observe DA transients during food approach, consistent with DA facilitating appetitive approach (34–38). The magnitude of these approach-associated transients decreases with successive pellets within a meal and recover at resumption of feeding after an intermeal interval. These data suggest that dopamine contributes to meal patterning, i.e., stopping and starting meals, by tracking short-term fluctuations in hunger and satiety. Earlier microdialysis studies demonstrate that extracellular dopamine increases during feeding (4,5) and FSCV studies demonstrate dopamine transients associated with appetitive approach (38–40). We extend these observations demonstrating subsecond DA transients associated with food approach are sensitive to short-term fluctuations in hunger and satiety that modulate meal patterning.

### Fasted mice: dissociation between meal initiation and termination

As expected, overnight fasting induced compensatory eating, reflected in both increased meal size (delayed meal termination) and meal frequency (increased meal initiation), but these effects were time-dependent across feeding. In the first hour, fasted mice initiated more meals and delayed meal terminations, resulting in larger, more frequent meals that substantially increased consumption. In the second and third hours, frequency of meal initiation remained elevated, but meals are *shorter* than the non-fasted mice, where shorter, more frequent meals result in reduced consumption in hours 2-3. This suggests a dissociation between the mechanisms regulating initiating and terminating meals, where the former reflects a persistent incentive-salience process that orients behavior toward food pursuit while the latter reflects a more variable satiation-feedback process that terminates food intake. DA transients on food approach were only altered (increased) in fasted mice during the first hour where they exhibited delayed meal termination and not in hours 2-3 where more frequent meals are combined with shorter, truncated meals. This suggests that modulation of feeding-related dopamine activity *during* feeding regulates whether the animal continues or terminates a meal.

One interpretation of these data is that the energy deficit generated by fasting induces a persistent motivation to seek food, i.e., enhanced incentive-salience of food-related stimuli, which leads to increased initiation of food pursuit and more frequent meals. However, this elevated food seeking is tempered by feedback from satiation. Specifically, compensatory eating during the first hour generates proportionally larger post-ingestive satiety signals. In hours 2-3, the fasted mice initiate more meals but operating in a context of greater post-ingestive satiety signals, meals are truncated sooner. Nevertheless, meal frequency and interest in food (incentive-salience) remains elevated in hours 2-3, fasted mice eat shorter but more frequent meals. Thus, increased initiation effectively compensates for increased satiety signals that arise from compensatory eating, while earlier termination prevents over-eating. Our data lacks identifiable behaviors during the intermeal interval that would allow us to assess alterations in dopamine activity related to decisions to initiate a meal.

### Pharmacological satiety decouples DA modulation from fluctuating hunger and satiety

Consistent with previous studies, Ex-4 decreased meal size (27,41). Despite a robust reduction in consumption, we observe no discernible effect of on mean amplitude of DA transients at pellet approach. However, Ex-4 abolished the modulation of these transients by hunger and satiety within and between meals as observed above. Ex-4 treated mice do not exhibit greater amplitude DA transients at the beginning of a meal that decline with successive pellets, nor larger transients at the start of meals following longer intermeal intervals as observed in non-treated mice. These data suggest that the modulation of dopamine that tracks short-term fluctuations in hunger and satiety are flattened out in Ex-4 treated mice.

In our data, DA transients on food approach in the Ex-4 treated mice seem always in a state analogous to the end of a meal. A recent report showing stimulation of DA neurons can overcome semaglutide induced satiety (24) is consistent with these observations. Our interpretation differs: rather than overcoming GLP1-induced satiety, we suggest that GLP-1 decouples DA from fluctuating hunger and satiety, i.e., unlinking a cardinal motivational effector normally responsive to changes in hunger and satiety, consistent with suggestions that GLP-1 agonists suppress the sensitizing effect of hunger on DA release (10).

Prior studies of GLP-1 agonists’ effects on mesolimbic dopamine have yielded inconsistent results. Fortin and Roitman (2017) observed no changes in DA in the NAc after Ex-4 administration. However, in the VTA, GLP-1 agonists decrease excitatory synaptic input onto mesolimbic DA neurons (42) and inhibit their burst firing activity (43). Our data is consistent with Fortin and Roitman (2017) in that we do not observe any decrease in phasic dopamine activity in the accumbens; however, we do observe the loss variability in those transients that would normally be linked to fluctuating hunger and satiety. This GLP-1 decoupling could be achieved by selectively disengaging inputs to the VTA that otherwise would carry hunger-satiety state information. As GLP-1 agonists are being explored in other compulsive behaviors, such as addiction (44), it would be interesting to further explore whether a *disengagement* of DA mediated motivational enhancement rather than simple decreases or increases in DA is a generalized effect of these agents.

### The timescales of dopamine, hunger and satiety

We observe minute-to-minute fluctuations in the magnitude of dopamine transients at food approach that correspond to short-term satiation and recovery of hunger across bouts of stopping and starting eating. This suggest that dopamine plays a role in the microstructure of feeding patterns, modulating *both* the drive to pursue food due to hunger and the diminution of that drive through satiation. Post-ingestive signals that modulate midbrain dopamine, such as insulin, ghrelin, glucose, provide a logical possible mechanism for linking dopamine to fluctuating hunger-satiety, but the changes we observe - e.g., declining dopamine transients with successive pellets occur at a timescale of less than a minute, i.e., 10s of seconds between pellets. A pellet cannot be ingested, travel the alimentary canal, be digested, absorbed and provide a post-ingestive signal rapidly enough to modulate the dopamine peak at the next pellet approach occurring ∼10 seconds later. Nevertheless, our data indicates dopamine does fluctuate on this timescale during active eating.

One mechanisms that could act at such a rapid timescale is vagal transmission. Fernandes et al (2020) demonstrate that gastric infusion of sucrose can rapidly increase burst activity in dopamine neurons. However, gastric infusion eliminates travel time down the esophagus and the breakdown of the pellets into glucose. Moreover, in Fernandes et al (2020), sucrose infusion *increased* dopamine activity to *facilitate* feeding, suggesting this rapid mechanism would increase, not decrease DA activity.

Satiation has multiple components that operate at different timescales, including early, rapid sensory and cognitive components and more delayed post-ingestive and post-absorptive signals (45–47). Sensory and cognitive components are essentially learned expectations that associate sensory information (chewing, taste and smell of pellet) with anticipated nutritional content based on prior experience. Sensory information during consumption may activate these learned associations and modulate dopamine signaling based on *expected* post-ingestive signals, providing a mechanism of fast satiation consistent with changes in dopamine from pellet to pellet within a meal.

When the signaling that mediates this anticipatory satiety during consumption is terminated after the animal stops eating, this allows an ’accounting’ as post-ingestive and post-absorptive signals come on-line that establish a new state of hunger-satiety. If this new state is less than full satiety, then ’hunger is recovered’ and eating resumes, repeating the process. As this cycle repeats, the slower timecourse post-ingestive/absorptive signals accumulate until they reflect a state of full satiety, and eating stops for an extended period. This process is analogous to the behavior of Agouti-related peptide (AgRP) neurons in the hypothalamus, whose activity signals hunger (48–51). AgRP neurons reduce their activity when food is available or expected to be available, before it is actually consumed, indicating that these neurons drive food *pursuit* but not consumption itself. If, however, the animal does not consume the encountered food, then AgRP activity returns to its previous higher level (52).

Acting on common downstream targets, competing neuropeptide signals that modulate satiation can be seen as discrete bursts of hunger - and satiety - promoting signals, such as neuropeptide Y (NPY) and alpha-melanocyte-stimulating hormone (α-MSH), respectively (53). Each signal can temporarily dominate or suppress the other based on changing metabolic state. During feeding, the balance gradually shifts with satiety-promoting signals becoming more frequent and sustained, while hunger-promoting signals are increasingly suppressed. This dynamic allows satiation to be adapted to on-going changes in energy ingestion and availability (53). This gradual emergence of satiety comprises many brief, stochastic, and competing peptide events (seconds) that are cumulatively integrated. Our data suggests that modulation of dopamine in relation to meal patterning may be linked to these competing, fluctuating signals as a mechanism of coupling on-going energy assessment with behavioral activation.

Importantly, although we see an *overall* pattern of decrease in DA transients associated with approach across meals, this is not a cleanly, monotonically decreasing function. Instead, variability is high; transient amplitude decreases only on average across a meal, but within a meal, individual DA transients associated with pellet approach may go up or down. This variability suggests the confluence of multiple processes. Three primary modulators might include: vagal stimulation and other signals that *increase* DA activity during consumption (54) to maintain on-going feeding, sensory-cognitive satiation as discussed above that constrains and pauses feeding, and the slower timecourse energy signals, including rising post-ingestive/absorptive signaling as well as pro-feeding signals, such as ghrelin that reflect newly updated energy states. All of these change at different timecourses each with its own variability, such that the net effect on DA at any given moment is highly variable. Despite this complexity and variability, we nevertheless can clearly discern a pattern of changing DA transients related to food approach that correspond to short-term fluctuations in hunger and satiety.

Finally, we often (informally) think of dopamine as *determining* behaviors. Our data highlights the fundamental neuromodulatory and stochastic nature of dopamine. Nevertheless, dopamine effects on behavior, noisy and stochastic as the signal may be, can be profound.

## Acknowledgments

This work was supported by a Klarman Family Foundation Eating Disorders grant (JAB), NIH National Institute on Drug Abuse grant DA052871 (JAB), the Spar Biosciences Laboratory (JAB), generously funded by Dr. Ira Spar, and by a PSC- CUNY Award, jointly funded by The Professional Staff Congress and The City University of New York (JAB).

## Author Contributions

JAB and DM developed the conceptual framework, designed the study, and wrote the manuscript. DM, CEM, EH, and KC collected the data. DM analyzed the data. FG mapped and analyzed histological data and did sensor validation. All the authors approved the final manuscript as submitted.

## Conflicts of Interest

No potential conflict of interest relevant to this article was reported.

## Notes

### Competing Interest Statement

The authors have declared no competing interest.

### Summary of Updates

Revised manuscript moved GCaMP recording from striatum to supplemental material, added a analysis of potential sex effects to supplementary material and generally revised the narrative of the results and discussion.

