## supplemental materials for "Dopamine signaling tracks naturalistic short-term fluctuations in hunger-satiety"

### Supplemental Material: GCaMP recordings

#### GCaMP recordings in the direct and indirect pathway in a free-feeding experiment with no food restriction or GLP-1

We compared four groups of mice distinguished by the population and location of cells from which we recorded GCaMP calcium signals (see methods): (1) dSPNs in the nucleus accumbens core (NAcc), (2) dSPNs in the nucleus accumbens shell (NAcSH), (3) iSPNs in the NAcc, and (4) iSPNs in the NAcSH, referred to as dSPN-core, dSPN-shell, iSPN-core, and iSPN-shell, respectively. Here, we report pathway-specific SPN calcium signals from all mice. Total consumption (pellets per day), the number of meals per day, and average meal size did not differ significantly across the defined groups of mice (suppl

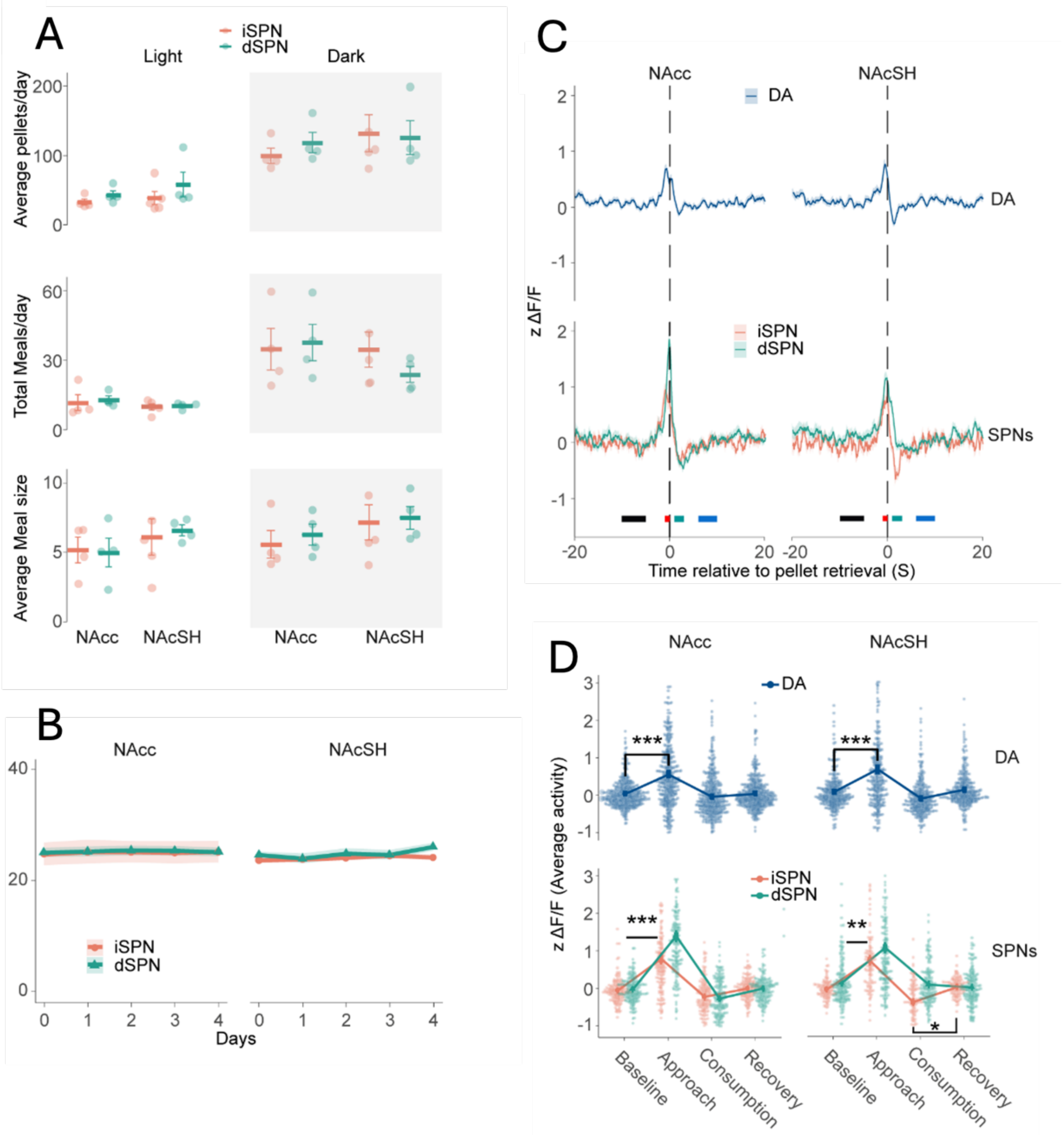


**Figure S1.** SPN activity increases during food approach in natural feeding. **A)** The total number of pellets, meals, and average meal size for all four days of feeding by groups and cycle. **B)** Body weights over four days by groups. **C)** Average dopamine release (top) and SPN Ca++ activity relative to pellet retrieval for direct and indirect SPN activity plotted by pathway and sub-region of the nucleus accumbens. Black, red, green, and blue rectangles indicate feeding stages: baseline, approach, consumption, and recovery, respectively. **D)** Average activity of DA and d/iSPNs during feeding stages for all groups. Data are presented as the mean ± s.e.m. in **p* < 0.05; ***p* < 0.01; ****p* < 0.001.

Fig 1A), nor did their body weights (Figure 1B). The mice exhibited consistent, stable feeding patterns across all groups and days.

SPN activity in both the direct and indirect pathways increased robustly during food approach immediately preceding pellet retrieval in both the NAcc and NAcSH (suppl Figure 1C-D, lower panels, $F_{(3, 2366)} = 341.8$, $p < .001$), consistent with prior reports (1). During consumption, a decrease was observed in the indirect pathway in the shell only (suppl Figure 1D, Tukey post-hoc: $t_{(2366)} = -4.14$, $p < 0.01$). These data indicate that transient increases in SPN activity correlates to food approach rather than consumption. These patterns of transient increase in SPN Ca++ signals mirror patterns observed in dopamine. We include the dopamine traces from the main paper (Fig 1I) broken out by the four groups for comparison.

*dSPNs activity attenuated as mice reached satiation in the Nacc*

As with dopamine, we evaluated progressive changes in SPN activity at pellet approach within and between meals. In both the NAcc and NAcSH, dSPN activity gradually decreased with successive pellet retrieval within a meal (suppl Figure 2A, green traces), an effect less clear in iSPNs (Figure 2A, orange traces). Comparing the first and last pellets of meals, we found in that both dSPN activity (suppl Figure 2B, Tukey: $t_{(880)} = 5.53$, $p < .001$) and iSPN activity (suppl Figure 2B, Tukey: $t_{(880)} = 3.85$, $p < .01$) significantly decreased on approach across pellets within a meal in the NAcc. In NAcSH, there was a trend toward reducing dSPN activity (suppl Figure 2B, Tukey: $t_{(880)} = 4.39$, $p = 0.079$), but no decrease was observed in iSPN activity (suppl Figure 2B, Tukey: $t_{(880)} = 0.69$, $p = .99$).

Examining changes in SPN activity between meals, dSPN activity increases from the last pellet of a preceding meal to the first pellet of the next meal in the NAcc (suppl Figure 2C, Tukey: $t_{(768)} = 4.65$, $p < .001$) but not in the NAcSH (suppl Figure 2C, Tukey: $t_{(768)} = 3.33$, $p = 0.20$). This suggests, in the NAcc, that the diminution of dSPN activity at pellet retrieval across successive pellets within a meal recovers between meals, similar to the recovery observed in DA. There is no significant recovery in iSPNs in the NAcc (suppl Figure 2C, Tukey: $t_{(768)} = 3.22$, $p = 0.26$) or NAcSH (suppl Figure 2C, Tukey: $t_{(768)} = 0.01$, $p = 0.99$).

To determine if a higher initial peak of SPN activity for the first pellet of a meal corresponds to longer meals, we compared d/iSPN activity from all meals larger than one pellet to single pellet meals across feeding stages. Consistent with DA release in the NAcc, we found that only dSPN activity during pellet approach is significantly higher in meals larger than a single pellet approach in the NAcc (suppl Figure 2D, top, dSPNs on approach, Tukey: $t_{(266)} = 3.97$, $p < 0.01$; iSPNs, Tukey: $t_{(260)} = 2.11$, $p = 0.41$) but not in the NAcSH (suppl Figure 2D, bottom, dSPNs on approach, Tukey: $t_{(266)} = 0.796$, $p = 0.99$).

We examined whether recovery of the initial peak in SPN activity at meal onset was influenced by the time elapsed since the last meal. In the NAcc, recovery of both dSPN and iSPN activity peaks on approach for the first pellet of the next meal was increased by the length of time since the last meal (suppl Figure 2E, dSPNs: $F_{(1, 57)} = 11.9$, $p < 0.001$; iSPNs: $F_{(1, 30)} = 6.74$, $p < 0.05$). In the NAcSH, only the dSPNs demonstrated a relationship between peak activity at first pellet in a meal and time elapsed (suppl Figure 2E, dSPNs: $F_{(1, 38)} = 6.88$, $p < 0.05$; iSPNs: $F_{(1, 51)} = 0.029$, $p = 0.865$). As with DA signals, time elapsed between meals had no effect on other behavioral segments, i.e., baseline, consumption, and recovery. To test whether DA and SPN activity correlate, we performed cross-correlations for each mouse. In all mice, DA correlated significantly with both dSPNs and iSPNs in the NAcc, both within and between meals, and within dSPNs but not iSPNs in the NAcSH (suppl Figure 3A-C).

Overall, dSPN activity mirrors DA activity, exhibiting clear modulation in signaling in the NAcc with regard to pellet approach and changes across and between meals. This dSPN eating related modulation is also evident in the NAcSH though less robustly. iSPN activity also shows eating related modulation, similar to DA and dSPNs, though only in the core and less robustly.

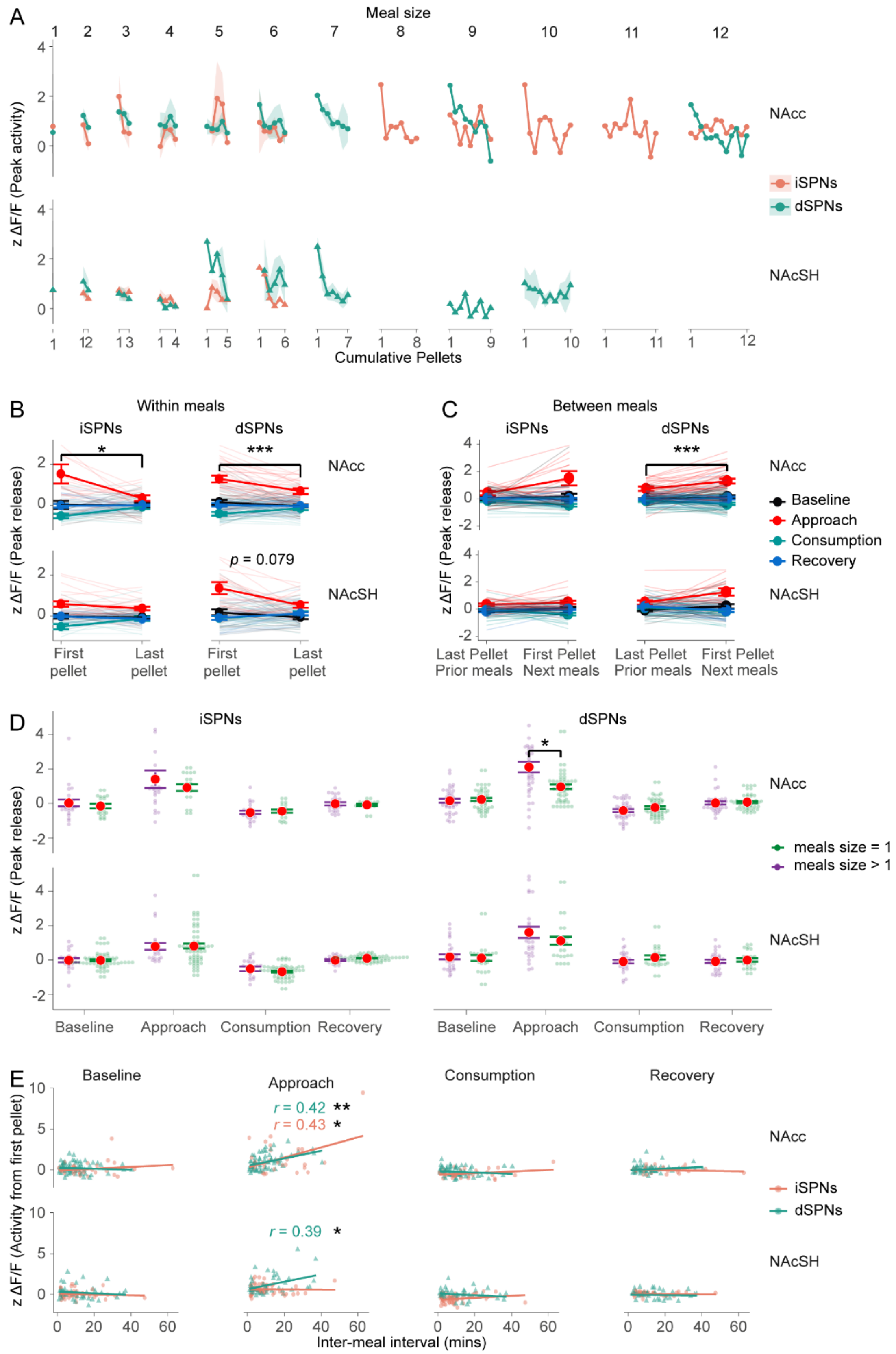


**Figure S2.** dSPNs activity attenuated as mice reached satiation in the NAcc**. A)** The average d/iSPNs activity in the NAcc and NAcSh was grouped by meal size across cumulative pellets during the approach. **B)** Average d/iSPNs activity from the first and last pellet of a meal grouped by the NAcc and NAcSH. **C)** Average d/iSPNs activity from the last (prior meal) and first (next meal) pellet between meals grouped by the NAcc and NAcSh. **D)** Average SPN activity from the last and first pellet between meals grouped by the NAcc and NAcSH. **E)** For each feed stage, a best-fit line is drawn for the inter-meal interval in relation to the d/iSPN activity. Activity is at the first pellet of a meal.

*Conclusions from SPN Ca++ signaling data*

Overall, we observe the similar patterns in direct and indirect pathway activity as we observed with DA. First, we see the same clear increase at pellet approach in both pathways. Ca++ transients that coincide with increased dopamine in the direct pathway are expected as DA activated the dopamine D1 receptors on these cells to promote dMSN activity. The mechanism for an increase in iMSN activity co-occurring with DA transients is less clear, given that iMSNs expression dopamine D2 receptors that are inhibitory. The lack of iMSN inhibition from increased dopamine might be accounted for the high affinity of D2 receptors already saturated at basal extracellular DA concentrations such that transient increases in DA yield no additional inhibition (2). We speculate the increased iMSN activity co-occurring with DA transients are coincident and not driven by dopamine. Instead, increased iMSN activity might be driven by increased glutamate or disinhibition from inhibitory interneurons. Our data do not speak to what mechanism mediates this concurrent increase in iMSN activity at pellet approach. However, this pattern of complementary d- and i- MSN activity-- in contrast to opponent processes-- has been widely observed and confirmed (3,4). Complementary increased iMSN activity may serve to strongly inhibit off-task activities, creating a higher bar for gating or releasing a specific behavior that must be driven by selective dMSN activity. The selected behavior could be exempted from iMSN inhibition via activity dependent corticostriatal LTD (5). The unique decrease in iSPN activity in the shell after consumption could reflects a brief period of diminished restrictiveness in corticostriatal selection to facilitate brief postprandial pause, a moment of possible exploration as the animal weighs pursuing another pellet or, possibly another activity.

After the initial experiment, we discontinued recording Ca++ signals from MSNs. On one hand, these data mirror the DA signals, leading to somewhat repetitive observations. On the other hand, the MSN activity is more difficult to interpret, offering multiple possible interpretations. Narrowing these interpretations would require additional experiments expanding the scope beyond our original intention. Thus, after this initial experiment, we focused only on DA for the fasting and GLP-1 studies. We report these GCaMP findings here for interested readers.

### Supplemental Material: Analysis of sex as a variable

*Rationale for supplemental sex analysis*: Our primary objective was to evaluate how dopamine signaling might correlate with progressive satiety and contribute to meal patterning. If we observe a role of DA in mediating satiety and meal patterning, a secondary question for future studies would be whether this dopamine contribution differs between sexes. The present study was not sufficiently powered to comprehensively evaluate sex differences.

Nevertheless, single sex studies are non-preferred, regardless of the goal of the study. Using both sexes, even if underpowered to serve as a study of sex differences, can provide an initial indication of whether there might be sex differences worth investigating in a larger subsequent study, once the core phenomenon is established. Here, we identify all statistically significant sex differences that emerged in our study (Table 1).

| Figure | Description | Significance | Interpretation |
|---|---|---|---|
| 1H-I | Peak DA for all events | *p* < .*001* interaction: sex x region | **(suppl Fig 1)** DA increase at approach reduced in males in NAcSH: no interpretation |
| 4D | # of meals | *p* < .*01* sex main effect | **(suppl Fig 2)** females initiated fewer meals (2A-B) that were (non-significantly) larger (2C); same total consumption (2C) |
| 4D | intermeal interval | *p* < .*05* sex main effect | see above |
| 5B | between meal DA recovery (fasting) | *p* < .*05* interaction: sex x recovery | **(suppl Fig 3)** females show less robust recovery following intermeal interval |
| 5E | correlation intermeal interval and peak at first pellet of next meal (fasting) | *p* < .*01* sex main effect | **(suppl Fig 4)** no interpretation, likely spurious |
| 7B | between meal DA recovery (GLP-1 agonist) | *p* < .*05* interaction: sex x hour | **(suppl Fig 5)** females show greater contrast between hours 1 and 2; no interaction with GLP-1 condition |

Table 1: Observed sex differences.

SEX DIFFERENCES OBSERVED

**Magnitude of DA transients across phases of behavior**

The DA transients across male and female mice were remarkably similar in baseline, approach, consumption, and recovery (suppl Fig 1), except for a reduced increase at pellet approach in the nucleus accumbens shell in males. We cannot speculate on whether this is a biologically meaningful difference or simply spurious. We note that differences in DA transients did not arise elsewhere in these studies, suggesting this is likely spurious. We cannot offer any potential interpretation.

**Meal patterning sex difference**

The remainder of the detected sex differences reflect differences in meal patterning. Females eat fewer meals, reflected in longer intermeal intervals (suppl Fig 2A-B). Though not statistically significant, presumably because of high variability, their meal size is likely larger, resulting in similar total consumption (supple Fig 2C-B).

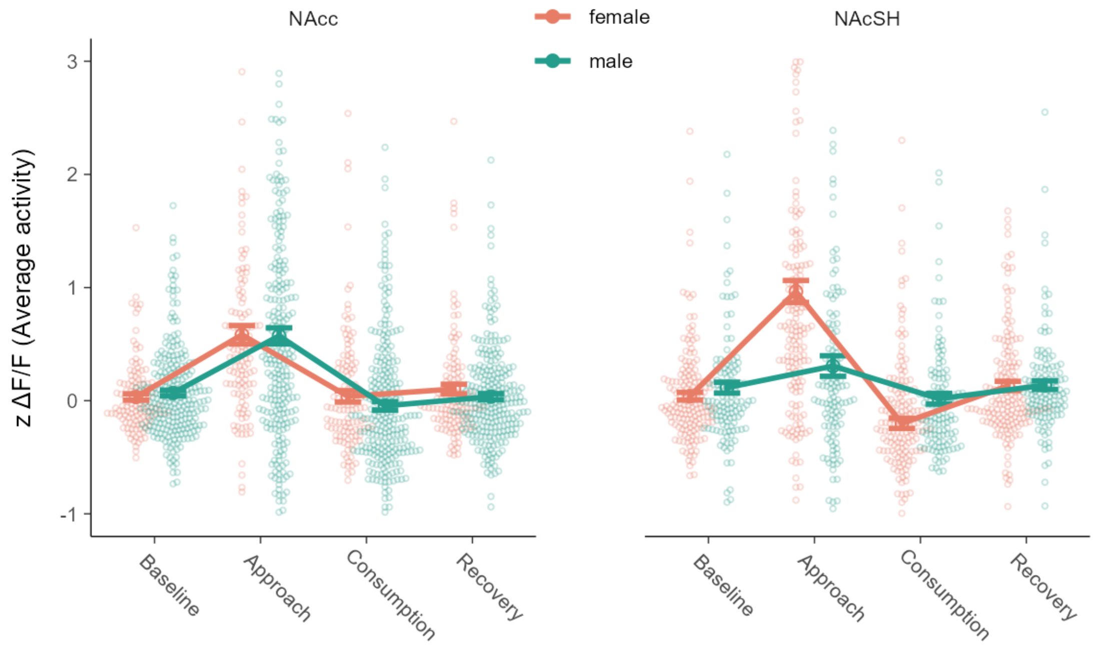


**Figure S3.** Dopamine transients at phases of eating by sex in the nucleus accumbens

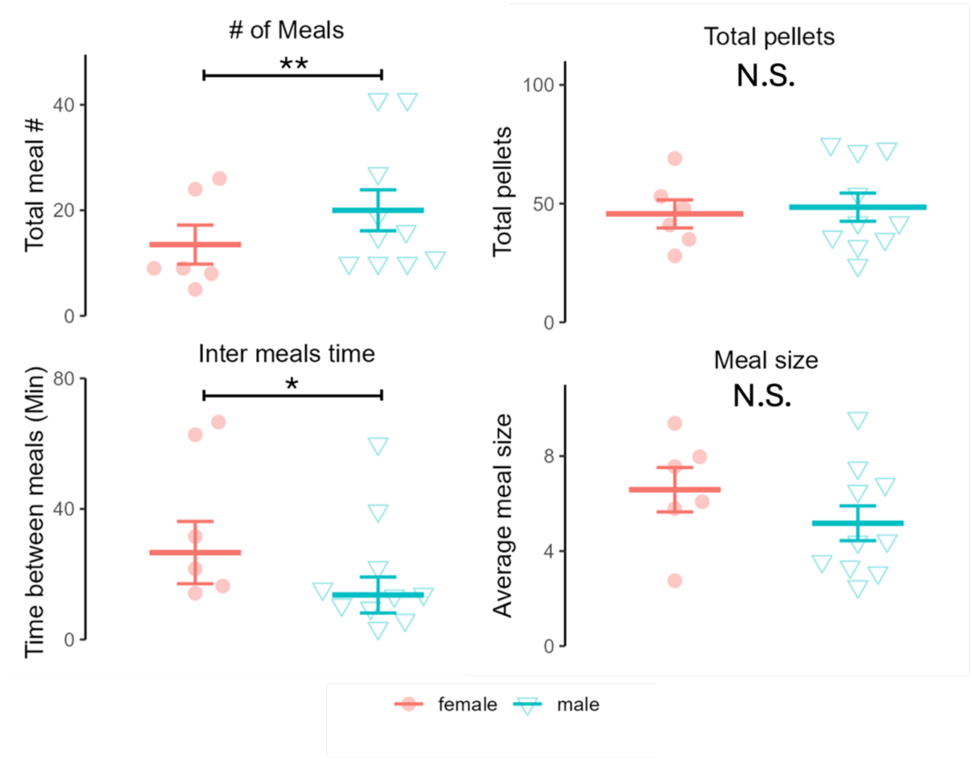


**Figure S4.** Meal patterning by sex.

In the fasting study, we observe a sex effect on the recovery of dopamine between meals, where females show a less pronounced recovery following an intermeal interval (suppl Fig 3), consistent with a sex difference in meal patterning. In the fasting experiment, there is a sex difference in the relationship between length of the intermeal interval and the subsequent DA peak at first pellet of the next meal (suppl Fig 4), though we find this uninterpretable.

Finally, in the GLP-1 agonist study: in Fig 7B in our report, there is no discernible difference between dopamine comparing across drug condition (Ex-4, vehicle) or hour (hour 1 vs. 2). When we break this out by sex (suppl Fig 5), we can see that there is a sex difference across hours; specifically, the males show very similar DA transients on approach in both hours while the females exhibit increased DA on approach in the second hour. This difference is observed across both drug conditions, suggesting it reflects an underlying difference in meal patterning across time. We cannot speculate on why the dopamine transient on approach might increase in the second hour in females as intuitively one would expect the opposite.

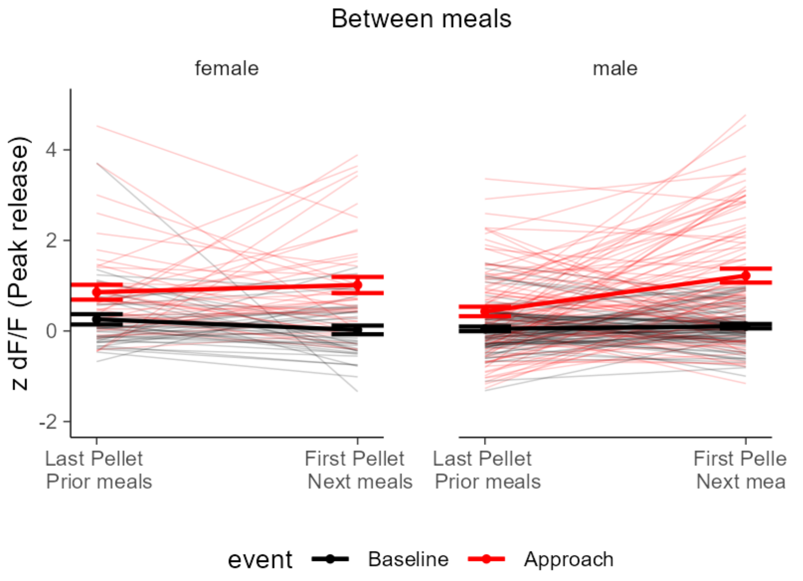


**Figure S5.** Recovery of DA transient magnitude at first pellet of next meal after an intermeal *interval*.

These analyses point to a potential difference in meal patterning between male and female mice, particularly in the regulation of meal number. We observed little difference in DA across the sexes and so cannot determine the degree to which DA might contribute to this potential meal patterning difference, which could arise from multiple possible sources.

Overall, while our conclusions must be qualified by low power for evaluating sex differences, we observed little differences between males and females. Where differences did emerge, these center around meal number and recovery of hunger following intermeal intervals. We observed no interactions between sex and the effects of either fasting or the GLP-1 agonist and little difference in DA signaling, suppl Fig 5 being the exception.

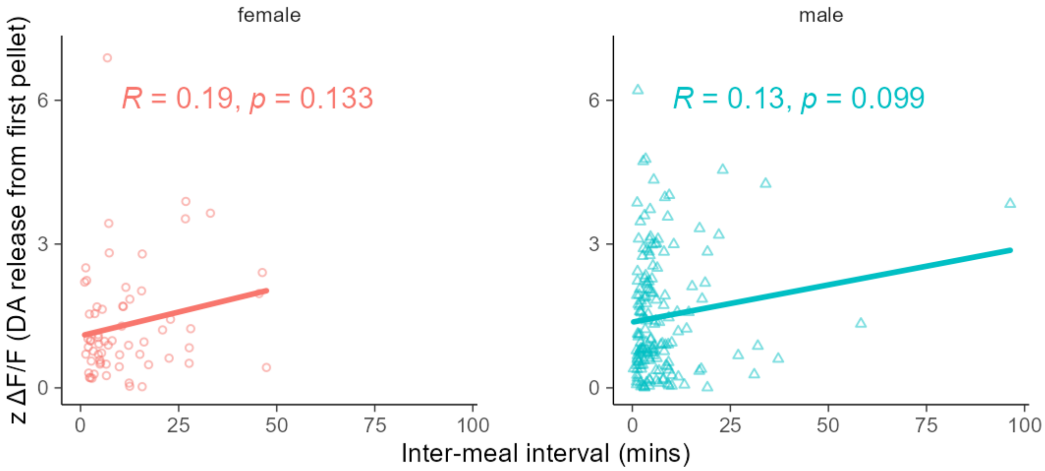


**Figure S6.** Relationship between magnitude of DA peak at first pellet of meal and length of intermeal interval.

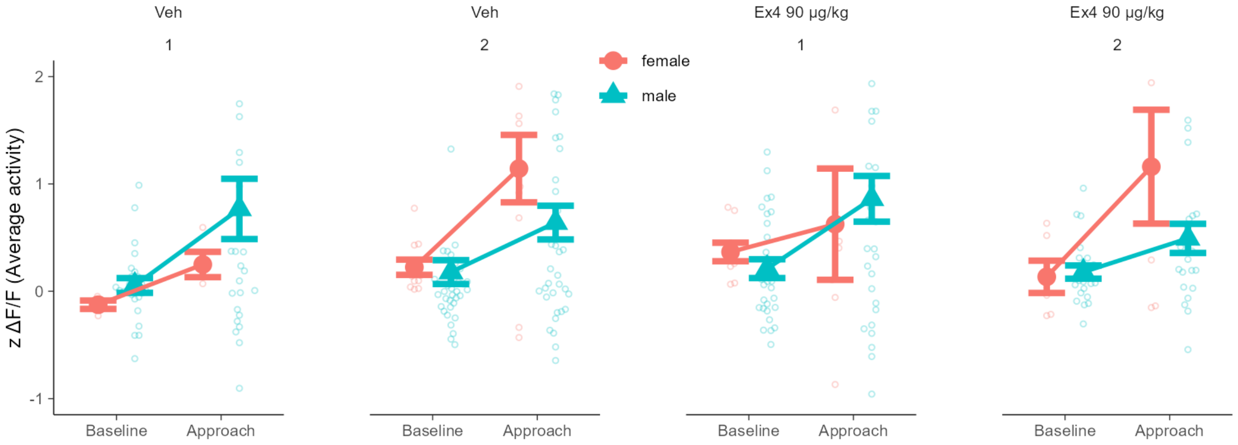


**Figure S7.** Increase in dopamine at approach by sex across hours and drug condition.

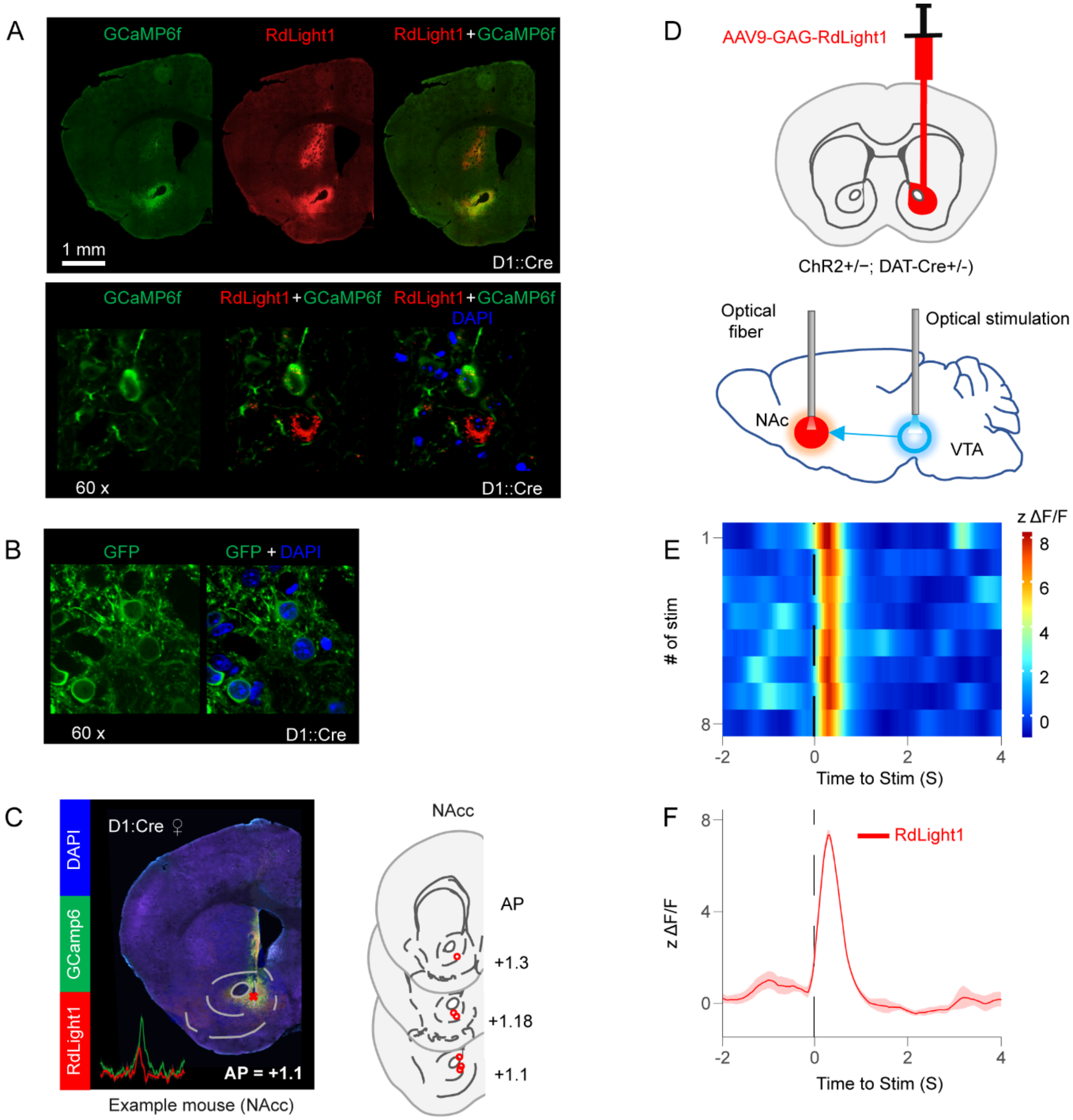


**Figure S8.** Histology and functional verification of recording preparation. **A)** For immunofluorescence, a D1: Cre mouse was virally injected with RdLight1 and GCaMP6f in the ventral striatum. Three weeks later, the mouse was anesthetized and then perfused, sectioned, and mounted. RdLight1 and GCaMP6f were immunostained using anti-RFP/ anti-GFP antibody, stained with DAPI, and imaged on confocal (top). Visual confirmation of GCaMP6f and RdLight expressing neurons (bottom). **B)** Numerous healthy GCaMP6f-expressing neurons. **C)** Example mouse of the histology verification of RdLight1 and GCaMP6f with the corresponding average DA (red) and SPN (green) traces (left). Fiber optic cannula (FOC) placement (right). **D)** Schematic of virally injected with RdLight1 in NAcc (top), optically stimulating VTA and recording DA release in the NAcc (bottom). **E)** Heatmaps DA release triggered by optical stimulation. **F)** Mean DA release (z scores), aligned to optical stimulation. n=8.
